# Development of a fluorescence reporter system to quantify transcriptional activity of endogenous p53 in living cells

**DOI:** 10.1101/2022.08.03.502578

**Authors:** Tatsuki Tsuruoka, Emiri Nakayama, Takuya Endo, Shingo Harashima, Rui Kamada, Kazuyasu Sakaguchi, Toshiaki Imagawa

## Abstract

The tumor suppressor p53 plays a central role in cellular stress responses by regulating transcription of multiple target genes. The temporal dynamics of p53 are thought to be important for its function: it encode input information and are decoded to induce distinct cellular phenotypes. However it remains unclear to what extent the temporal dynamics of p53 reflects the activity of p53-induced gene expression. In this study, we report a multiplexed reporter system that allows us to visualize the transcriptional activity of p53 at the single cell level. Our reporter system features simple and sensitive observation of the transcriptional activity of endogenous p53 to the response elements of various target genes. Using this system, we show that the transcriptional activation of p53 exhibits strong cell-to-cell heterogeneity. The transcriptional activation of p53 by etoposide is highly dependent on the cell cycle but not by UV-C. Finally, we show that our reporter system allows simultaneous visualization of the transcriptional activity of p53 and cell cycle. Our reporter system can thus be a useful tool for studying biological processes involving the p53 signaling pathway.

**Summary statement:** Tumor suppressor protein p53 is one of the hub factor of the cellular stress responses. Our novel reporter system allows us to visualize the transcriptional activity of endogenous p53 easily and precisely.

## Introduction

The tumor suppressor protein p53 functions as a hub protein for a variety of signaling pathways involved in cellular stress responses (Bieging and Attardi, 2012; Kastenhuber and Lowe, 2017). The *TP53* gene, which encodes p53, is the most frequently mutated gene in many human malignant tumors, suggesting the potential importance of p53 in suppressing tumorigenesis (Olivier et al., 2010). p53 is known to act as a transcriptional regulator, and to be activated by a variety of post-translational modifications induced by various cellular stresses, including hypoxia, nutrient starvation, and genotoxic stresses such as DNA damage. It is well known that the tetramerization of p53 is essential for p53 activation (Kamada et al., 2016). Activated p53 recognizes and binds to a large number of p53 response elements (*p53RE*s) in the genome, leading to the transcriptional regulation of multiple target genes. In this way, p53 inhibits tumorigenesis by inducing various cellular responses such as apoptosis, cell cycle arrest, and DNA repair (Appella and Anderson, 2001; Horn and Vousden, 2007; Meek, 2009). Previous reports suggested that there are more than 300 target genes of p53 (Fischer, 2017).

In recent years, it has become clear that temporal dynamics plays an important role in intracellular signal transduction (Purvis and Lahav, 2013). In the p53 pathway, the temporal dynamics of p53 is known to play a pivotal role in the regulation of p53-regulated cellular functions. Temporal dynamics of p53 differed depending on the type of cellular stress, inducing distinct cellular responses (Chen et al., 2013; Purvis et al., 2012; Yang et al., 2018). It has also been shown that different doses of the same type of stress induce different temporal dynamics of p53 and consequently lead to different cellular phenotypes (Chen et al., 2013; Paek et al., 2016; Yang et al., 2018). Similar temporal regulation was also observed in other signaling pathways, including the ERK, NF-κB, and NFAT pathways (Lee et al., 2009; Marshall, 1995; Nelson et al., 2004; Yissachar et al., 2013). Recently, it has been proposed that cells encode input cellular information into the temporal dynamics of signaling molecules and then decode this information into downstream pathways (Purvis and Lahav, 2013). These mechanisms are thought to play a pivotal role in transmitting cellular information efficiently. However, it has been reported that different cellular responses are induced even though the artificially induced temporal dynamics of p53 expression have the same shape as the natural temporal dynamics (Purvis et al., 2012). This suggests that the expression level of p53 does not necessarily reflect its transcriptional activation level (Loewer et al., 2010). In fact, previous studies suggested that p53 has a very large number of post-translational modifications that regulate its stability, oligomerization status, and binding affinity for DNA and transcriptional cofactors in a complex and diverse manner (Murray-Zmijewski et al., 2008). Therefore, to understand the decoding process in the downstream of p53 signaling pathway, it is highly desirable to develop a system that can visualize the activation status of p53 instead of its expression level in single living cells.

In previous attempts to visualize the downstream of p53 signaling pathway in single living cells, transcriptional assay systems using *p53RE* and endogenous tags based on genome editing technology have been reported (Harton et al., 2019; Stewart-Ornstein and Lahav, 2016). The system using tagged target gene products is able to capture actual changes in protein expression. On the other hand, in terms of monitoring the decoding process of signaling factors, transcription reporters have the advantage of directly observing the transcriptional activity in a manner independent of mRNA stability, protein stability, and the translation rate of the protein. The reporter systems that can evaluate p53 tetramer levels, an activated species of p53, have also been reported (Gaglia and Lahav, 2014; Gaglia et al., 2013).

In this study, we report a multiplexed reporter system that enables monitoring of the transcriptional activity of endogenous p53 at the single cell level with high sensitivity. This reporter system is based on the *p53RE*-based transcriptional activity assay system previously reported in yeast-based functional assay (Kato et al., 2003; Shimada et al., 1999). Our reporter system is characterized by an internal standard for the transcriptional activity of p53, and by replacing the *p53RE*, the difference in the *p53RE* in each target gene can be easily and quantitatively evaluated. In addition, by tracking the fluorescence in the nucleus, it is possible to easily observe the temporal change of transcriptional activity for each single cell. Analysis using our reporter system revealed that the transcriptional activity of p53 was highly heterogeneous even among genetically identical cells. We also found that the transcriptional activation of p53 under etoposide treatment is cell-cycle dependent and shows stronger cell-cycle dependency than the p53 protein level.. Finally, we demonstrate that the transcriptional activity of p53 can be visualized simultaneously with other cellular events when used in combination with other reporter systems.

## Results

### Development of a multiplexed reporter system

We developed a novel reporter system to quantitatively analyze the transcriptional activity of endogenous p53 in individual living cells (Fig. 1A and Fig. S1A,B). This reporter system is based on the underlying mechanism by which p53 recognizes a specific p53 response element (*p53RE*) in the p53 target gene and regulates its transcription. This reporter system enables us to quantify the activation status of p53 by the expression unit of (*p53RE(C3)*)-Cerulean-NLS and the p53-independent constitutive expression unit of (*simian virus 40 promoter (SV40)*)-mCherry-NLS, respectively. The reporter system can also monitor the transcriptional activity of p53 against the endogenous *p53RE* derived from the *CDKN1A* gene by the expression unit of the (*p53RE(CDKN1A)*)-Venus-NLS. We call these reporters (*C3)*-Cerulean, (*SV40)*-mCherry, and (*CDKN1A)*-Venus, respectively. Since a NLS is attached to the C-terminus of the fluorescent protein, the expression level of each reporter can be evaluated by quantifying the fluorescence intensity in the nucleus. These reporter genes were knocked into the *AAVS1* site of human lung adenocarcinoma cell line A549 (p53 wild type) by sequence-specific cleavage with CRISPR/Cas9 and site-specific recombination. The *AAVS1* site was selected as a knock-in site because it is an open chromatin region and is less susceptible to repression of gene expression (Sadelain et al., 2011). Another advantage of using the *AAVS1* site is that the copy number of the reporter gene is expected to be the same in all cells. These features of the *AAVS1* site allowed us to quantitatively capture the transcriptional activity of p53. The fluorescent proteins selected for inclusion in the reporters are suitable for precisely monitoring transcriptional activity because of their high brightness and rapid chromophore formation (Nagai et al., 2002; Rizzo et al., 2004; Shaner et al., 2004). The fluorescence spectra of the three fluorescent proteins do not overlap in the observed wavelength range, so there are no effects from leakage into the other fluorescent channels.

**Figure 1.**
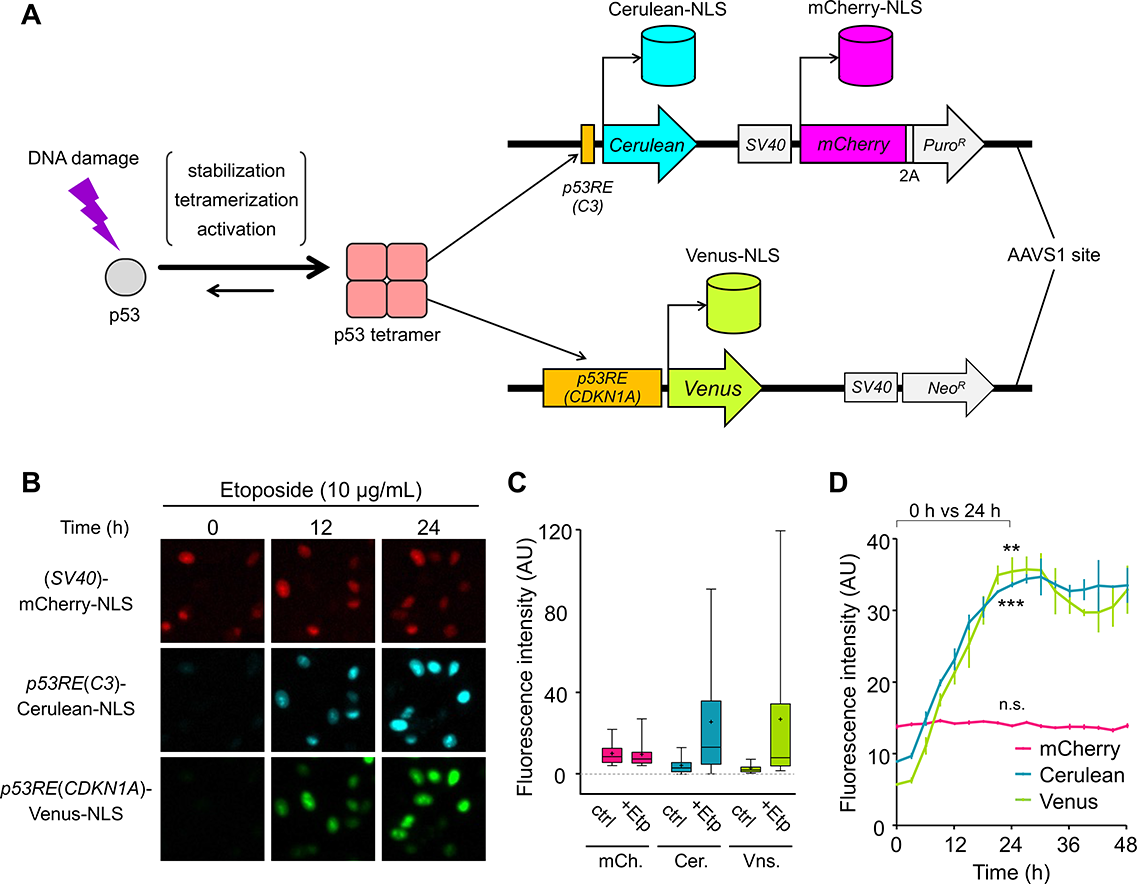
Development of the fluorescence reporter system for monitoring the transcriptional activity of endogenous p53. A) Schematic representation of the reporter system. B) Microscope images of the reporter cell line derived from A549 cells (p53 wild type) after the addition of 10 μg/ml etoposide. Scale bar: 50 μm C) Distribution of the fluorescence intensity at 0 h (left) and 24 h (right) after 10 μg/ml etoposide treatment. Horizontal lines and crosses indicate the medians and means of distributions, respectively. Boxes and whiskers include the values between the 25th and 75th and the 5th and 95th percentiles, respectively. More than 2,500 cells were analyzed under each condition. D) Temporal changes of the fluorescence intensity after the addition of 10 μg/ml etoposide. The plot shows the mean fluorescence intensity and S.D. from three independent experiments. More than 500 cells were used for each experiment. Statistical analysis was performed by using the fluorescence intensity at 0 h and 24 h. n.s.= not significant (p > 0.05). **p < 0.005, ***p < 0.0001, unpaired t-test.

We characterized the properties of the novel reporter system as follows. Etoposide was used as a genotoxic stressor because etoposide induces DNA double-strand breaks (DSBs) by inhibiting topoisomerase II, which activates the p53-mediated DNA damage responses (Nitiss, 2009a; Yang et al., 2018). When the reporter cells were treated with etoposide, the fluorescence intensity of *(C3)*-Cerulean and *(CDKN1A)*-Venus, both of which represent p53-dependent transcriptional activity, significantly increased, while the fluorescence intensity of *(SV40)*-mCherry, which represents p53-independent transcriptional activity, did not change (Fig. 1B). The fluorescence intensity of *(C3)*-Cerulean and *(CDKN1A)*-Venus showed large distribution after the etoposide treatment, whereas the *(SV40)*-mCherry intensity was almost unchanged (Fig. 1C and Fig. S1C). This result indicates that the transcriptional activation of p53 in response to genotoxic stress is heterogeneous even among genetically identical cells. Time-lapse imaging of the reporter cells showed that the fluorescence intensity of *(C3)*-Cerulean and *(CDKN1A)*-Venus increased from 6 h after stimulation, reaching a peak at 27–30 h, while the intensity of *(SV40)*-mCherry did not change at any time point (Fig. 1D). Analysis with the reporter system showed that the fluorescence intensity of (*C3*)-Cerulean and (*CDKN1A*)-Venus increased in a dose-dependent manner with etoposide concentration (Fig. S1D). We found that there were differences in dose dependence between (*C3*)-Cerulean and (*CDKN1A*)-Venus. These differences may result from transcriptional regulation of the *CDKN1A* gene by other factors or differences in affinity between p53 and the p53-binding site (Gartel and Radhakrishnan, 2005; Weinberg et al., 2005). The fluorescence intensity of *(SV40)*-mCherry did not change under all dose conditions tested. We also verified the p53-dependent response of the reporter system; a p53-knockout reporter cell line, which was confirmed by genomic PCR and Western blotting (Fig. S1E,F), showed only slight changes in the fluorescence intensity of *(C3)*-Cerulean and *(CDKN1A)*-Venus following etoposide treatment (Fig. S1G). Taking these results together, we concluded that this reporter system specifically monitors the transcriptional activity of p53. It is unclear why a slight transcriptional activation was observed in the p53-knockout cell line, but this may have been due to the other p53-family proteins, p63 and p73; both of these proteins share considerable amino acid sequence identity with p53 and are known to control the expression of p53-regulated genes (Levrero et al., 2000).

### Transcriptional activation of six different p53 target gene reporter cell lines

A notable feature of our reporter system is that an artificial p53 response element, *p53RE(C3)* is used separately from the endogenous *p53RE* to serve as an internal standard for p53 transcriptional activity. Therefore, by replacing the endogenous *p53RE* with that of another target gene, differences in transcriptional regulation between target genes can be quantitatively analyzed by comparison with this internal standard. To demonstrate this, a total of six reporter cell lines (including *CDKN1A*) were established using *p53RE* regions derived from different target genes (*CDKN1A*, *BAX*, *MDM2*, *GADD45*, *RRM2B*, *14-3-3σ*). After 24 h of etoposide treatment, these reporter cell lines showed no significant change in the distribution of *(SV40)*-mCherry fluorescence intensity, which exhibits a constitutive expression level (Fig. 2A). On the other hand, the distribution of *(C3)*-Cerulean fluorescence intensity, an internal standard for p53 activation status, increased in all cell lines, but the distribution did not show any significant differences (Fig. 2B). In contrast, the distribution of *p53RE (each gene)*-Venus fluorescence intensity, which exhibits transcriptional activity against the endogenous *p53RE*, showed a large variation among the different cell lines, with particularly large heterogeneity in transcriptional activity against *p53RE (CDKN1A)* (Fig. 2C). Normalization of the transcriptional activity for each *p53RE* with the internal standard showed that some genes differed greatly in distribution while others did not (Fig. 2D). In particular, the *CDKN1A* gene exhibited strong transcriptional activation under our experimental condition, followed by *14-3-3σ* gene. The other four genes showed a similar distribution, but with a slight increase in activity for *BAX* and a slight decrease in activity for *RRM2B*. Scatter plots of the transcriptional activity of each *p53RE* relative to the internal standard showed differences in this relationship among the genes (Fig. S2). Specifically, for *p53RE(CDKN1A)* and *p53RE(14-3-3σ)*, the degree of increase in transcription activity from the endogenous *p53RE* relative to the internal standard showed a linear relationship, whereas, for the other genes, transcription activity tended to be rapidly increased when the internal standard exceeded a certain level. These differences in activation properties among *p53RE*s may be one means by which cells respond to cellular stress efficiently using specific hub factors.

**Figure 2.**
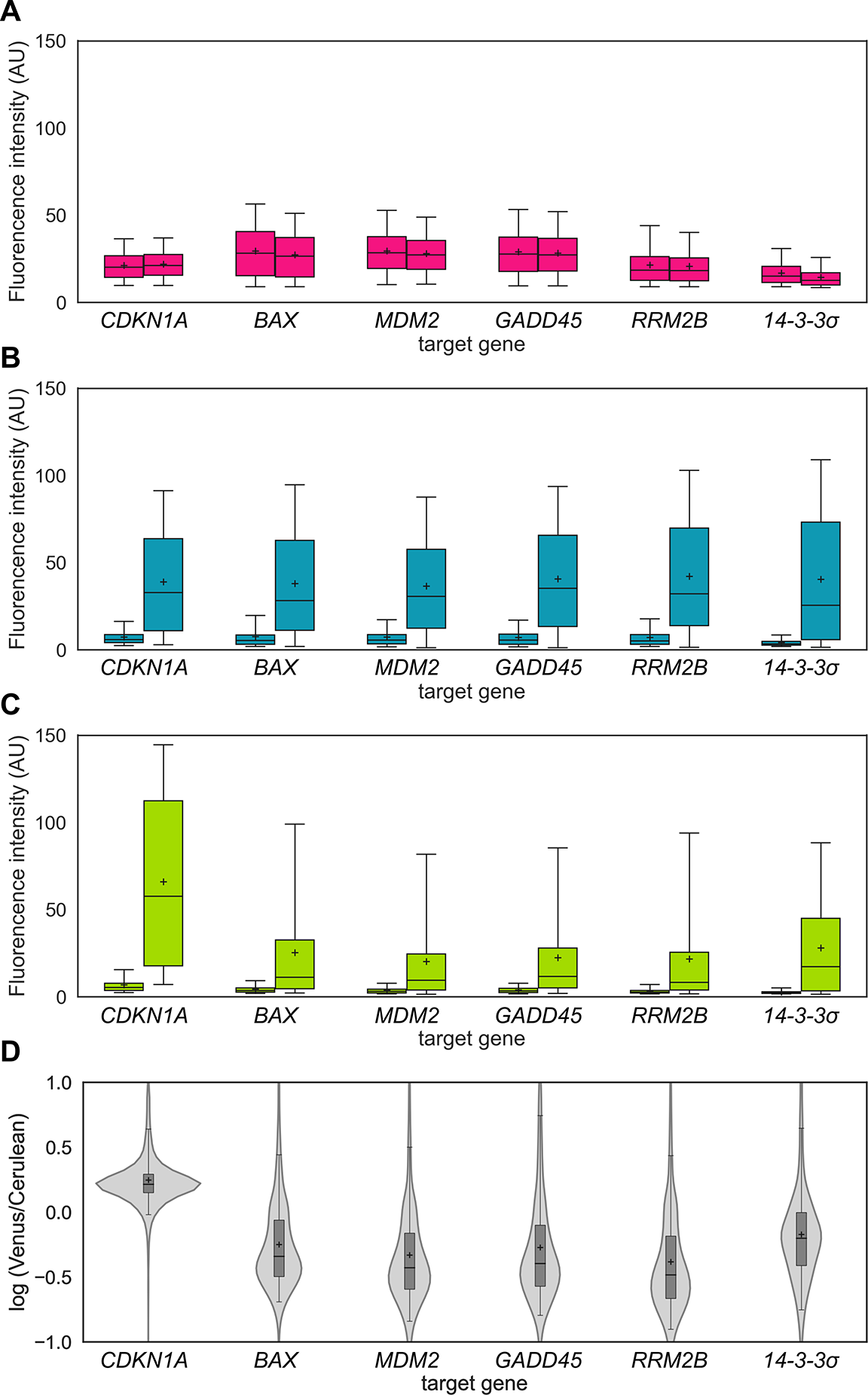
Transcriptional activation of six different p53 target gene reporter cell lines. A)-C) Distribution of the fluorescence intensity at 0 h (left) and 24 h (right) of the mCherry (A), Cerulean (B), and Venus (C) in reporter cell lines with *p53REs* derived from six different target genes after 10 μg/ml etoposide treatment. Horizontal lines and crosses indicate the medians and means of distributions, respectively. Boxes and whiskers include the values between the 25th and 75th and the 5th and 95th percentiles, respectively. More than 4,000 cells were analyzed under each condition. D) Distribution of Venus / Cerulean fluorescence intensity ratio of individual cell at 24 h after 10 μg/ml etoposide treatment for each reporter cell line. Horizontal lines and crosses indicate the medians and means of distributions, respectively. Boxes and whiskers include the values between the 25th and 75th and the 5th and 95th percentiles, respectively. Outliers (data with Venus / Cerulean ratios greater than 10 or less than 0.1) are excluded.

### Tracking the transcriptional activity of endogenous p53 in a single living cell

We analyzed how the transcriptional activation pattern of p53 differs from cell to cell when the same stress is applied to a homogenous cell population. The transcriptional activity of p53 was quantified simultaneously with detection of cell death by time-lapse live cell imaging. All subsequent experiments were performed using the reporter cell line with the *p53RE* region of *CDKN1A* because transcriptional activation for endogenous *p53RE* was most heterogeneous in this cell line. The reporter cells were manually tracked, and cell death was determined by cellular morphological changes (Fig. 3A and Fig. S3A; the images in Fig. 3A are related with that in Fig. 3B, cell trace 3). Interestingly, the pattern of p53 transcriptional activation was heterogeneous even within homogeneous cell populations (Fig. 3B). In addition, even between daughter cells derived from the same parental cell, the p53 activation pattern and the cellular response were often different (Fig. 3B). We next classified all cell traces into surviving cells and dead cells, and found that p53 transcriptional activation occurred significantly in cells undergoing cell death (Fig. 3C,D; data for *(SV40)*-mCherry are shown in Fig. S3B,C). Of note, some of the cell populations that did not undergo cell death also showed strong p53 transcriptional activation. We also confirmed that the p53-dependent transcriptional activity had little correlation with the constitutive expression level as measured by *(SV40)*-mCherry, indicating that the basal expression level is unlikely to be related to the transcriptional activity of p53 (Fig. S3D). These results strongly suggest that transcriptional activation of p53 is necessary for the induction of cell death. However, there was considerable overlap in the transcriptional activation of p53 between the cells that caused cell death and those that did not. This overlap again suggests a key role not only for the p53 signaling pathway but also for other signaling pathways involved in cell death by genotoxic stress, as previously reported (Roos et al., 2016).

**Figure 3.**
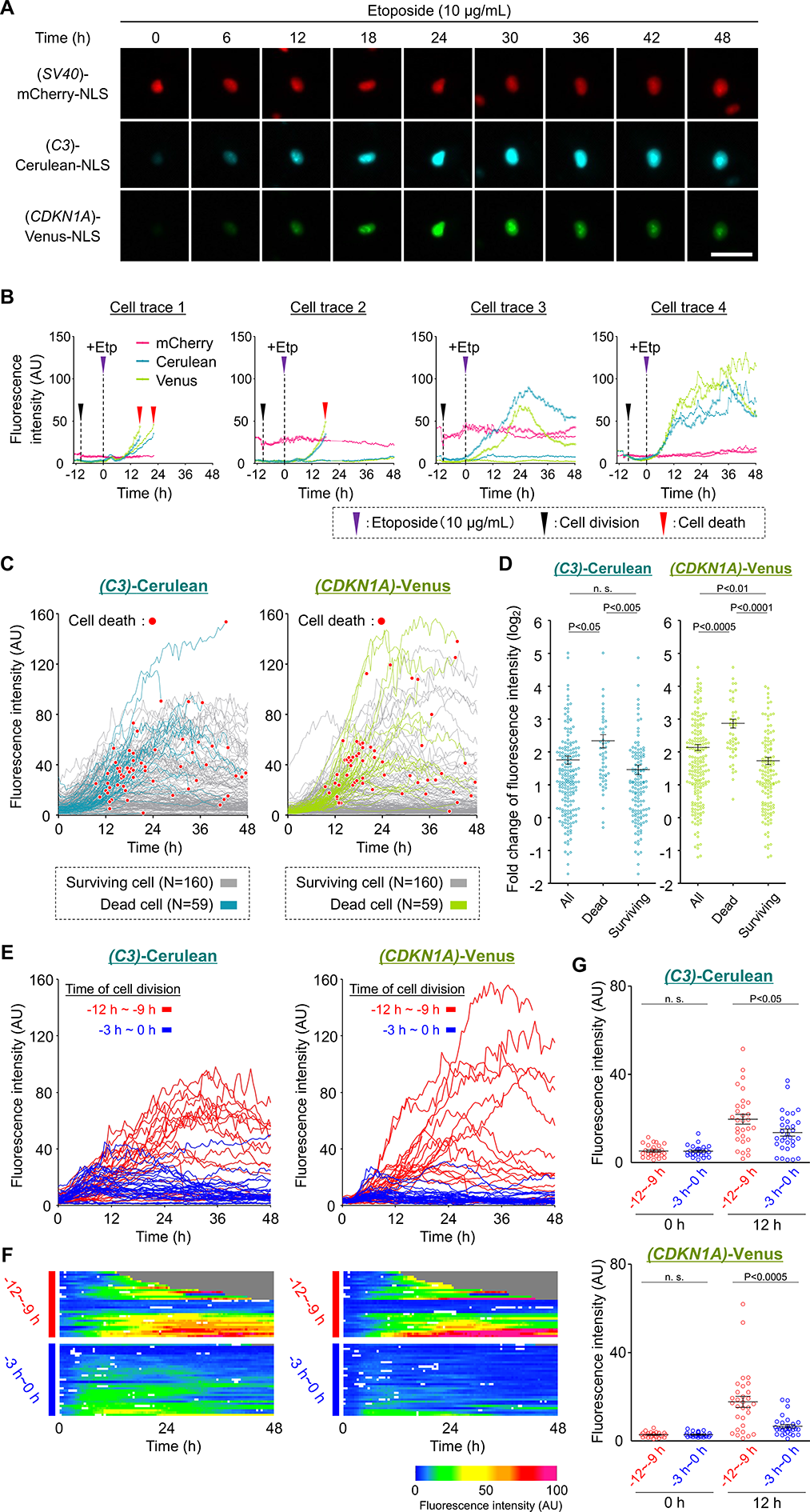
Quantification of the transcriptional activity of endogenous p53 at the single cell level. A) Time-lapse microscope images of the reporter cell line after 10 μg/ml etoposide treatment. Scale bar: 50 μm. B) Representative single cell traces of fluorescence intensity. C) Single cell traces (Cerulean, Venus) of all cells. Red dots indicate cell death. The gray or colored lines show the traces of surviving cells (n=160) and dead cells (n=59), respectively. D) Fold changes of fluorescence intensity (Cerulean, Venus) at 12 h compared with that at 0 h after 10 μg/ml etoposide treatment. Horizontal lines indicate the mean with S.E.M. P values were calculated by an unpaired t-test with Welch’s correction. n.s.=not significant (p > 0.05). E) Single cell traces classified by the time of cell division before 10 μg/ml etoposide treatment. Cells that underwent cell division 12–9 h (n=30) and 3–0 h (n=33) before etoposide treatment are shown in red lines and blue lines, respectively. F) Heatmaps of the single cell traces shown in (E). Gray traces indicate cells that had undergone cell death. White areas indicate that the cells were not traced. G) Distribution of fluorescence intensity at 0 h and 12 h after 10 μg/ml etoposide treatment. Horizontal lines indicate the mean with S.E.M. P values were calculated by an unpaired t-test with Welch’s correction. n.s.= not significant (p >0.05).

We classified all cell traces according to cell division time and found that the transcriptional activity of p53 was remarkably reduced in cells that divided 0 ~ 3 h before stress addition (presumably early G1 phase) compared to cells that divided 9 ~ 12 h before stress addition (presumably late G1 ~ S phase) (Fig. 3E; all data classified by cell division time are shown in Fig. S4A-C). In addition, the ratio of cell death decreased from 43% to 3% (Fig. 3F, gray area), respectively for these two groups. These results suggest that transcriptional activation of p53 by etoposide is dependent on the phase of the cell cycle, and thereby causes heterogeneous cellular responses. We also compared the fluorescence intensity before and after the stress addition and found no differences between the two groups before stress addition (Fig. 3G). This result indicates that the different patterns of transcriptional activation were not due to the initial activation state of p53. A previous report showed that the major cytotoxic effects of etoposide are exerted mainly in the S/G2 phase, and our observation is consistent with this report (Hainsworth and Greco, 1995).

### Transcriptional activation of p53 under UV-C stress condition

We also analyzed the p53 transcriptional activation patterns when cells were treated with UV-C (25 J/m^2^) irradiation and found that there was considerable cell-to-cell heterogeneity as well as under etoposide treatment (Fig. 4A). On the other hand, there was no clear correlation between the intensity of p53 transcriptional activation and cell death as in the case of etoposide treatment. Interestingly, comparison of the time courses of the mean values of surviving and dead cells revealed that p53 activation was stronger in surviving cells at an early stage after the stress addition (Fig. 4A, upper small panel, and Fig 4B). This result differs from that of etoposide treatment, indicating that the p53 transcriptional activation patterns differ depending on the type of stress.

**Figure 4.**
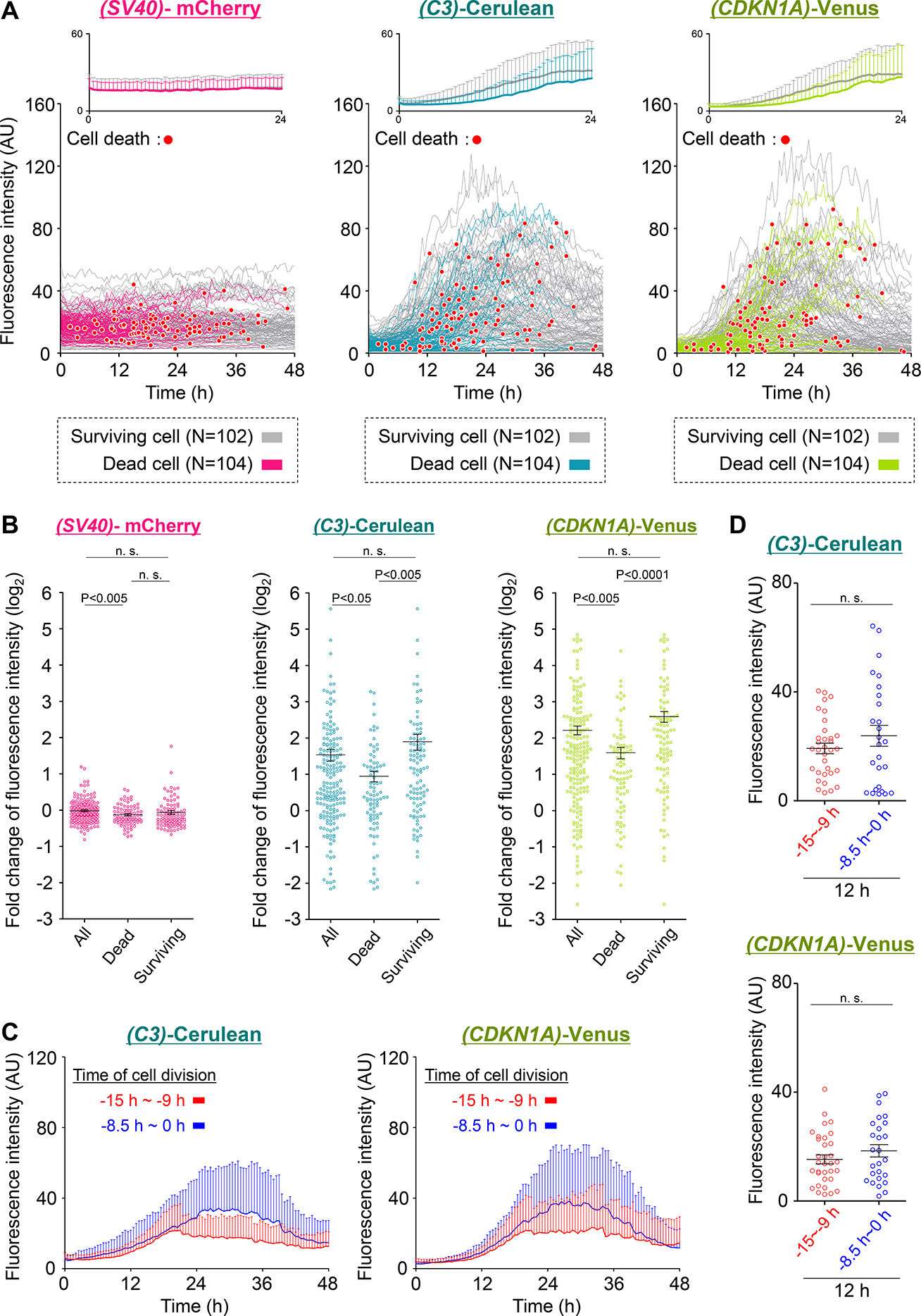
Quantification of the transcriptional activity of endogenous p53 after UV irradiation. A) Single cell traces of all cells. Red dots indicate cell death. The gray or colored lines show the traces of surviving cells (n=102) and dead cells (n=104), respectively. The upper small panel shows the mean + SD of the all cell trace from 0-24 h. B) Fold changes of fluorescence intensity at 12 h compared with 0 h. Horizontal lines indicate the mean with S.E.M. P values were calculated by an unpaired t-test with Welch’s correction. n.s.=not significant (p > 0.05). C) Average + SD plot of the single cell traces classified by the time of cell division before UV-C irradiation. Cells that underwent cell division 15–9 h (n=36) and 8.5–0 h (n=28) are shown in red and blue lines, respectively. D) Distribution of fluorescence intensity at 12 h after 25 J/m^2^ UV-C irradiation. Horizontal lines indicate the mean with S.E.M. P values were calculated by an unpaired t-test with Welch’s correction. n.s.= not significant (p >0.05).

We similarly analyzed the cell cycle dependency of p53 transcriptional activation patterns under UV-C irradiation condition. Comparison of p53 transcriptional activation patterns of cells dividing 0~8.5 h before stress addition (presumably G1 phase) or 9~15 h before stress addition (presumably S phase) showed no significant differences until around 20 h after stress addition (Fig. 4C,D). Thereafter, p53 transcriptional activity in the presumably S phase cells plateaued, whereas in the presumably G1 phase cells, transcriptional activity continued to increase until about 26 h after stress addition and then decreased. Interestingly, the ratio of cell death was 54% (15/28) for presumably G1 phase cells and 69% (25/36) for presumably S phase cells, indicating that the cell groups in which transcriptional activation of p53 occurred more strongly resulted in less cell death. Although details require further study, these results may indicate the importance of the effect of early p53 transcriptional activation on cellular responses under UV stress and the contribution of a p53-independent cell death-inducing pathway.

### Effect of the cell cycle phase for transcriptional activation of p53

We next examined the effect of the cell cycle on the transcriptional activation of p53 in more detail using the reporter cell line. The cells were synchronized in a specific phase of the cell cycle, double thymidine block (DTB) for G1/S phase synchronization and serum starvation (SS) for G1 phase synchronization. The cells were also synchronized at the G2/M phase by release 8 hours after DTB (Fig. 5A). We confirmed that the cell cycle was successfully synchronized under both conditions (Fig. S5A,B). The reporter cell line was synchronized and then treated with etoposide (Fig. 5B). After 24 h of etoposide treatment, the fluorescence intensity of G1-synchronized cells decreased 0.6-fold for *(C3)*-Cerulean and 0.5-fold for *(CDKN1A)*-Venus compared to that of asynchronous cells. In contrast, the fluorescence intensity of S-synchronized cells increased 1.7-fold for *(C3)*-Cerulean and 2.2-fold for *(CDKN1A)*-Venus. G2-synchronized cells did not show a significant change in transcriptional activity of p53. The level of *(SV40)*-mCherry was slightly decreased in synchronized cells. Although the details are not clear, it has been reported that CDK1 regulates the balance of the overall cell proliferation rate and global protein synthesis rate (Haneke et al., 2020). Therefore, the decrease in *(SV40)*-mCherry may be the result of a change in the protein synthesis rate linked to the cell proliferation rate. We also confirmed the effect of cell cycle synchronization on the p53 signaling pathway. The treatment of SS did not affect the transcriptional activity of p53 under our condition (Fig. S5C). The treatment of DTB induced slight activation of p53, but there were no differences in the fluorescence intensity of *(C3)*-Cerulean or *(CDKN1A)*-Venus at the time point of drug treatment (Fig. S5D). Hence, we considered that this difference would not have a significant impact on our experiments.

**Figure 5.**
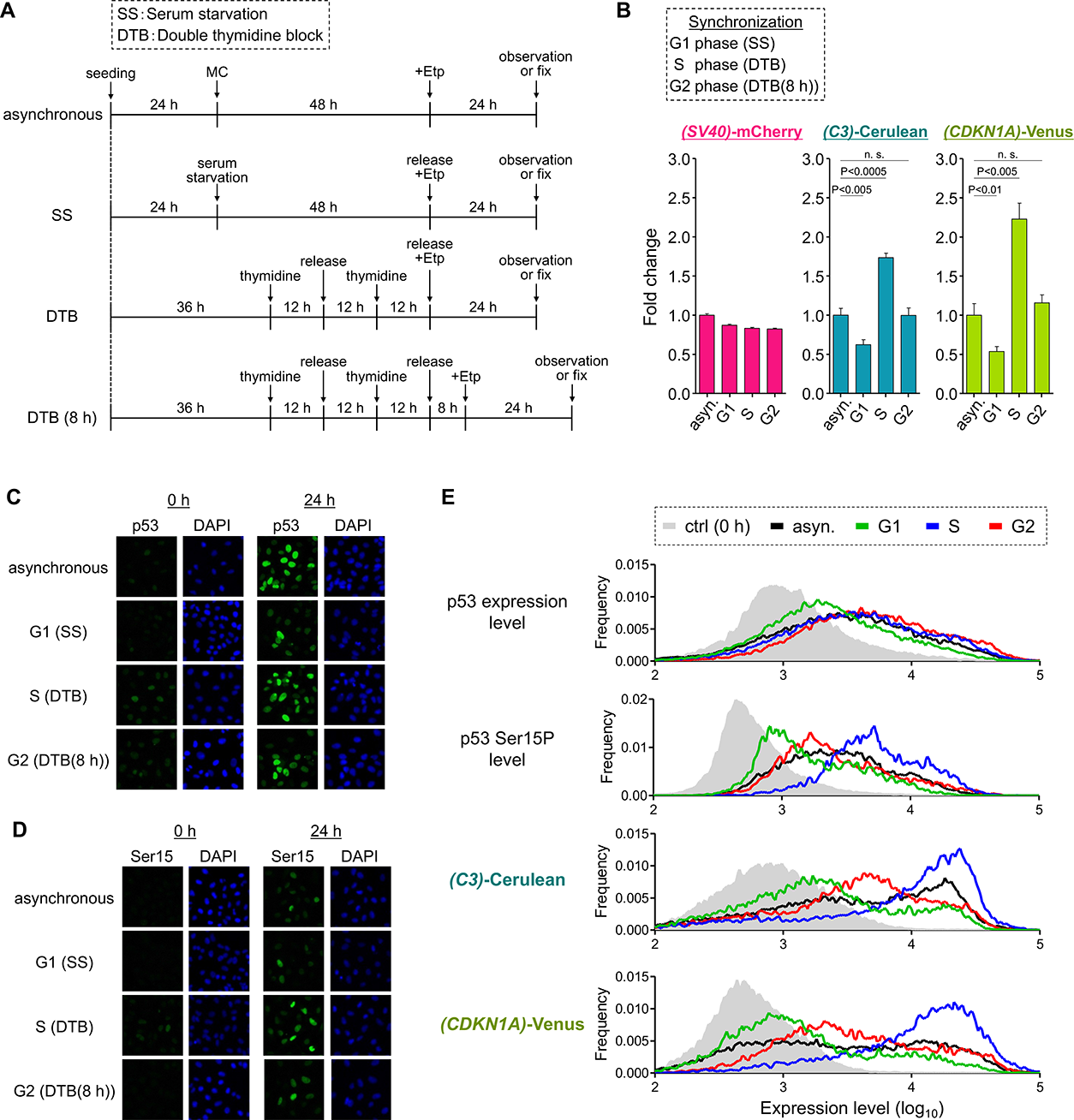
Effect of the cell cycle phase on the transcriptional activity of endogenous p53. A) Experimental outlines of the cell cycle-specific treatment by cell cycle synchronization. Cells were synchronized at the G1 phase by serum starvation and S phase by double thymidine block (thymidine concentration: 2 mM). Cells were also synchronized at G2 by culturing 8 h after removing thymidine. B) Fluorescence intensity at 24 h after 10 μg/ml etoposide treatment. Horizontal lines indicate the means with S.D. from three independent experiments (more than 500 cells were used in each experiment). P values were calculated by an unpaired t-test. n.s.= not significant (p > 0.05). C) p53 expression level at 0 h (left panel) and 24 h (right panel) after 10 μg/ml etoposide treatment. In each panel, left and right images show the immunofluorescence for p53 and nuclear area by DAPI staining, respectively. D) p53 Ser15P level at 0 h (left panel) and 24 h (right panel) after 10 μg/ml etoposide treatment. In each panel, left and right images show immunofluorescence for p53 Ser15P and nuclear area by DAPI staining, respectively. E) Distribution of the total fluorescence intensity after 10 μg/ml etoposide treatment. The panels show results of the p53 expression level and p53 Ser15P level quantified by immunofluorescence using normal A549 cells, *(C3)*-Cerulean intensity and *(CDKN1A)*-Venus intensity quantified by A549 reporter cells.

Several reports have shown that the expression level of p53 does not reflect its transcriptional activity (Loewer et al., 2010; Loffreda et al., 2017). Therefore, we analyzed the expression level of p53 and the level of p53 phosphorylated at Ser15 (p53S15P) after etoposide treatment by immunofluorescence staining using normal A549 cells (Fig. 5C,D). The obtained images were quantified and compared the distribution of fluorescence intensity (Fig. 5E). The expression levels of both p53 and p53S15P were increased under all conditions, but there were some differences among the conditions. Interestingly, the distribution of the p53 expression level in the S/G2-synchronized cells was almost the same as that in asynchronous cells, but the distribution of the p53 expression decreased in the G1-synchronized cells. On the other hand, the levels of the p53S15P were substantially increased in S-synchronized cells, but decreased in G1/G2-synchronized cells. The shift in the distribution of p53S15P was more significant in G1-synchronized cells, and was only slight in G2-synchronized cells. The distribution of the *(C3)*-Cerulean and *(CDKN1A)*-Venus also showed a shift to a higher expression level in S-synchronized cells and a lower expression level in G1-synchronized cells. The distributions of the *(C3)*-Cerulean and *(CDKN1A)*-Venus in G2-synchronized cells indicated an intermediate expression level (Fig. 5E). These results once again indicate that the expression level of p53 does not necessarily reflect the activation state of p53. Furthermore, the coefficient of variation of p53 expression levels and p53S15P levels calculated from the data under cell cycle-unsynchronized conditions revealed that the variation in p53 expression levels did not change significantly before and after the stress addition, while the variation in p53S15P levels significantly increased (Fig. S5E). This result may suggest that the heterogeneity in phosphorylated p53 levels is partly responsible for the cell-to-cell heterogeneity in p53 transcriptional activation by etoposide treatment. Our reporter system can directly quantify the transcriptional activity of p53 and may be a useful tool in studies involving the p53 signaling pathway.

### Simultaneous monitoring of cell cycle and transcriptional activity of p53 in living cells

Finally, we demonstrated the simultaneous imaging of p53-dependent transcription and cell cycle progression at the single cell level. We developed a novel reporter stable cell line using the Fucci reporter system for visualization of the cell cycle (Fig. 6A and Fig. S6A). Due to the limitation of available fluorescence channels, we used only the Cdt1 (30-120) fragment, which is stabilized in G1 phase and degraded in S/G2/M phase (Sakaue-Sawano et al., 2008). Hence, this reporter system monitors G1 phase-specific fluorescence by the Fucci-G1-Cerulean cassette, transcriptional activity against *p53RE (CDKN1A)* by Venus, and the constant expression level by mCherry. Time-lapse imaging of the reporter cells showed that the fluorescence intensity of Fucci-G1-Cerulean increases after cell division (Fig. 6B, white arrowhead; the images in Fig. 6B are related with that in Fig. 6D, cell trace 4). The fluorescence intensity of *(CDKN1A)*-Venus increased after etoposide treatment and decreased after removing the drug. To automatically classify these cell traces, we determined the classification rule (Fig. 6C; see the *Materials and methods* for detail). Using this rule, we classified all cell traces as early G1, late G1, S, and G2 by the cell cycle phase at drug treatment (Fig. 6C). We referred to these groups of cell traces as early G1, late G1, S, and G2-population, respectively. Applying this classification rule to the cell traces of this reporter system, we can simultaneously observe p53-dependent transcriptional activity, the cell cycle phase, and cellular response (Fig. 6D). All cell traces were then classified according to the classification rule and found that the transcriptional activation pattern of p53 had different characteristics that varied in a cell cycle-dependent manner (Fig. 6E; data for *(SV40)*-mCherry are shown in Fig. S6D). On average, the transcriptional activity of p53 in the G1-population was lower than that in S/G2-population. The ratio of cell death in the G1-population was also lower than that in S-population. In late G1-population, some cells showed high transcriptional activity of p53 and cell death. This was probably because the late G1 population contained the boundary population of G1/S phase cells. These cells showed on average high p53 transcriptional activity as well as a high ratio of cell death. Interestingly, the ratio of cell death in S-population was remarkably different from that in G2-population even though these two populations showed similar p53-activation levels. Transcriptional activation of p53 in G2-population occurred more rapidly than that in S-population and stopped at 12 h after drug treatment. These differences in activation patterns may have affected the final cellular outcome. We note that the pattern of Fucci-G1-Cerulean after drug treatment differed between G1 and S/G2-population. It is well known that etoposide induces mainly G2-arrest, and therefore, the pattern of Fucci-G1-Cerulean indicates that G1-population enter the S phase, and then all population are arrested at the G2 phase after drug treatment. Moreover, we confirmed that after release from etoposide cells failed to show a clear cell cycle re-entry over the next 12 h. Overall, our system proved to be useful for visualizing the transcriptional activity of p53 along with other biological phenomena.

**Figure 6.**
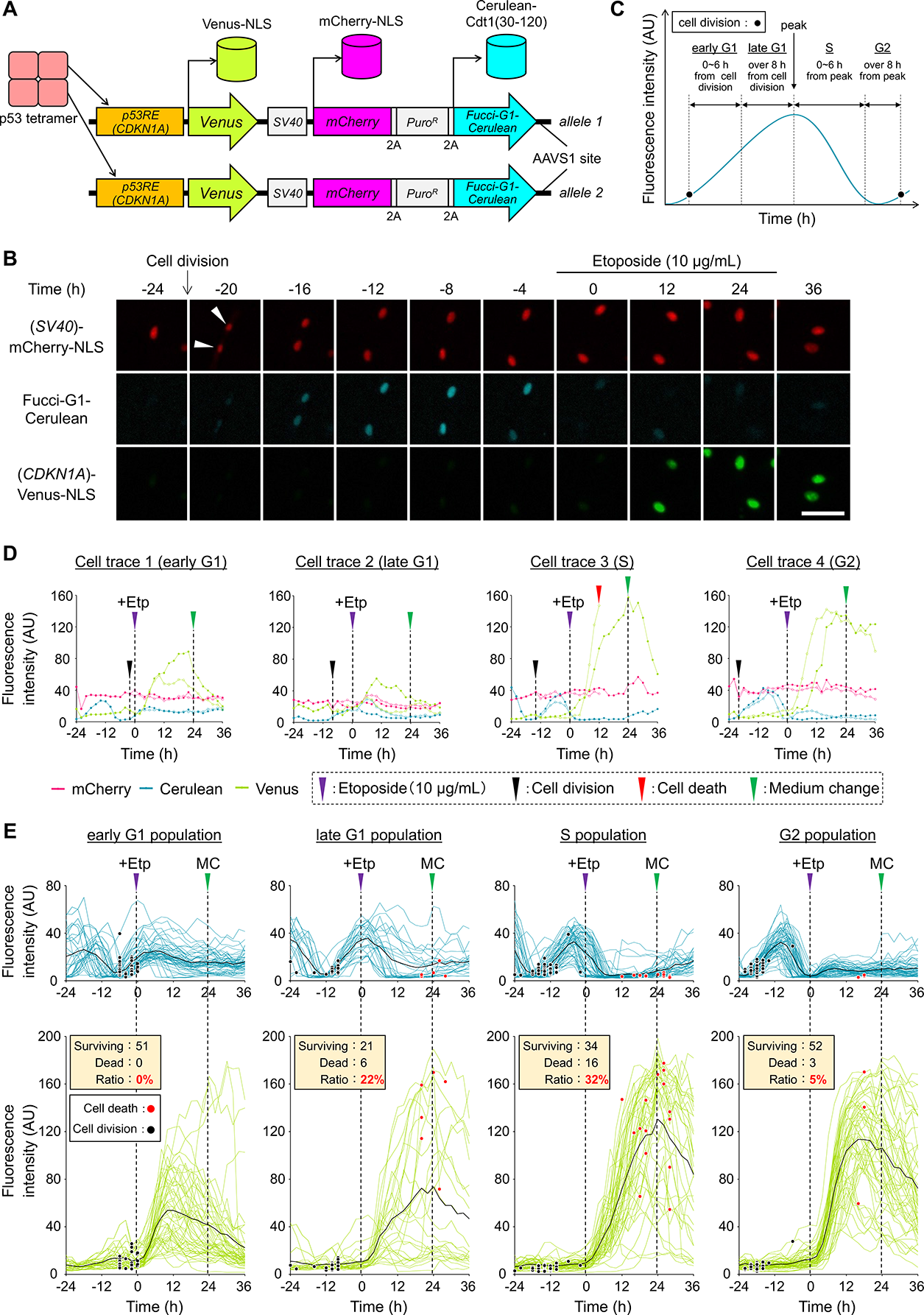
Simultaneous monitoring of the transcriptional activity of endogenous p53 and the cell cycle phase. A) Schematic representation of the reporter system used to quantify the transcriptional activity of p53 and the cell cycle. B) Time-lapse microscope images of the reporter cell line. Scale bar: 50 μm. C) Methods used to distinguish the cell cycle phase from the time course data of Fucci-G1-Cerulean. D) Representative single cell traces of fluorescence intensity. E) Single cell traces (upper panel: Cerulean, lower panel: Venus) of all cells classified by the cell cycle phase at the time point of stress addition. Black and red dots indicate cell division and cell death, respectively. The black line is a trace of the average value.

## Discussion

In this study, we have developed a reporter system that allows sensitive observation of p53 transcriptional activity in single living cells. Our reporter system is unique in that it uses *p53RE(C3)*, which consists only of p53 binding sequences, as an internal standard for p53 transcriptional activity. By comparing the responses to endogenous *p53RE* of several target genes with the responses to *p53RE(C3)*, it is possible to quantitatively analyze differences in transcriptional regulation among target genes. In fact, transcriptional activity for *p53RE(C3)* showed a similar distribution among all cell lines, while transcriptional activity for endogenous *p53RE* showed a different distribution among cell lines (Figure 2B,C). This result suggests that the responsiveness to each *p53RE* is affected by other factors such as p53 affinity, other regulators that bind to *p53RE*, and post-translational modifications of p53. We also calculated the ratio of Venus to Cerulean fluorescence intensity per cell to more clearly demonstrate differences in the responsiveness of endogenous *p53RE*s (Fig. 2D and Fig. S2). Our reporter system design integrating internal standards provides an useful framework for quantitative analysis of differences in responsiveness of multiple target genes regulated by hub transcription factors, not just p53.

We found that the transcriptional activation pattern of p53 under etoposide treatment differs among genetically identical cells (Fig. 3). It was previously reported that the temporal dynamics of p53 in A549 cells under etoposide treatment showed either “periodic pulsing” or “monotonic induction,” with the ratio of cells displaying “monotonic induction” increasing in a dose-dependent manner (Chen et al., 2013; Yang et al., 2018). It has also been reported that the majority of “periodically pulsing” cells exhibit cell cycle arrest, whereas the majority of “monotonic induction” cells result in cell death. In our study, although we did not observe the p53 expression and p53 transcriptional activity at the same time, the characteristics of the transcriptional activation patterns in the cells that caused cell death suggest that cells that showed strong p53 transcriptional activation corresponded with “monotonic induction” cells and cells that did not show strong p53 transcriptional activation corresponded with “periodic pulsing” cells. Previous studies have shown that *p53RE*s found in different p53 target genes had varying affinities for p53 and thus different thresholds for p53 expression levels (Harton et al., 2019; Wu et al., 2017). These differences may result in the separate decoding of a single temporal dynamics of p53 into transcriptional dynamics for each target gene. In addition, it is generally known that the affinity of p53 for *p53RE*s of genes associated with cell death is lower than that for *p53RE*s of genes associated with cell cycle arrest (Weinberg et al., 2005). These properties may be a part of the mechanisms that amplify differences in the temporal dynamics of p53 among cells and confer robustness to the signaling pathway.

We also showed that the pattern of transcriptional activation of p53 following treatment with etoposide is highly dependent on phase of the cell cycle. It is well known that the cellular response to etoposide is cell-cycle dependent, and that etoposide is the most cytotoxic during the S/G2 phase (Hainsworth and Greco, 1995). Our results are consistent with these conclusions, as the transcriptional activity of p53 tended to be higher in S/G2 cells than in G1 cells (Fig. 5B and 6E). The main reason for this cell-cycle dependency may be the difference in the total amount of DNA damage. Topoisomerase IIα, the primary cellular target of etoposide, is mainly responsible for resolving the topological problems associated with replication and chromosome segregation in the S/G2 phase (Nitiss, 2009b). In addition, topoisomerase IIα is known to be upregulated during the S/G2 phase (Goswami et al., 1996; Hsiang et al., 1988). Therefore, the amount of DNA damage caused by the same dose is expected to be higher in S/G2 cells. In fact, several reports showed a significant increase of DSBs in S/G2 cells (Muslimović et al., 2009; Tammaro et al., 2013). The cell-cycle dependency of p53S15P levels is probably closely related to these changes upstream of the p53 signaling pathway. On the other hand, transcriptional activation of p53 following UV-C irradiation showed a heterogenous response similar to that of etoposide treatment even though it was largely independent of the cell cycle. Interestingly, there was also little correlation between transcriptional activity of p53 and cell death events. It has been reported that under UV irradiation, cross-inhibition of c-Jun N-terminal kinase (JNK) via p38 generates cell-to-cell variability in JNK activity, resulting in stochastic cell death (Miura et al., 2018). It is also known that JNK has multiple phosphorylation sites on the transactivation domain of p53 and regulates p53 stability and transcriptional activity (Buschmann et al., 2001; Oleinik et al., 2007; Saha et al., 2012). Therefore, under UV-C irradiation, the heterogeneity observed in p53 transcriptional activity may be due to heterogeneity in upstream JNK activity, which may contribute to stochastic cell death. Although the detailed upstream mechanisms of the p53 signaling pathway under various stress conditions require further investigation, it is likely that heterogeneity in the p53 signaling pathway occurs at different layers due to various factors such as those mentioned so far. In addition, it has been reported that cisplatin, an anticancer drug which, like etoposide, is widely used in clinical practice, also shows almost no cell-cycle dependency, and these different characteristics between drugs are clinically important (Granada et al., 2020).

In our experiments, we did not find marked cell-cycle dependency in the expression level of p53 (Fig. 5C,E). On the other hand, the level of p53S15P showed a remarkable cell-cycle dependency, which is consistent with the results observed for transcriptional activity (Fig. 5D,E). A possible mechanism to explain this observation is the contribution of direct transcriptional repression of p53 by MDM2. MDM2 functions as an E3 ubiquitin ligase and regulates p53 expression, while MDM2 also directly represses transcription by concealing the transcriptional activation domain of p53 (Karni-Schmidt et al., 2016; Oliner et al., 1993). Therefore, the phosphorylation of Ser15 may disassociate p53 from MDM2 and increase the proportion of p53 that is free of MDM2, resulting in cell cycle-dependent transcriptional activation. MDM4, a homolog of MDM2 that does not have E3 ubiquitin ligase activity, but has direct transcriptional repressive activity, may also be involved in this mechanism (Karni-Schmidt et al., 2016). Additionally, inhibition of p53 tetramer formation by an apoptosis repressor with the caspase recruitment domain (ARC) may also be involved in the cell cycle-dependent response. ARC is known to be repressed by transcription through p53 and degradation by MDM2 (Foo et al., 2007; Li et al., 2008). Therefore, the formation of p53 tetramers is inhibited in a cell cycle-dependent manner. These factors would allow cells to regulate the transcriptional activity of p53 in a cell cycle-dependent manner without any marked changes in p53 expression levels.

Although p53 was strongly transcriptionally activated in both the S and G2 phases compared to the G1 phase, almost none of the cells in G2 phase underwent cell death (Fig. 6E). It has been reported that activation of p53 in the G2 phase is sufficient to induce cellular senescence (Johmura et al., 2014; Krenning et al., 2014). Therefore, most of the cells stressed in the G2 phase are thought to undergo cellular senescence. It is known that p21 plays an important role in the induction of cellular senescence. p21 is rapidly degraded in S-phase cells, which may induce a different response of the G2 phase by predominantly inducing the expression of cell death-related gene products (Bornstein et al., 2003). These findings can only be obtained by comparing global regulation at the transcriptional level with the expression of individual target gene products.

In this study, we developed an experimental system that enables us to monitor the activation status of p53 in single living cells. We have also demonstrated that the cell cycle is strongly involved in the heterogeneous response to genotoxic stress in genetically identical cells by single-cell tracking using this system. Our reporter system can be an important tool for providing new insights into the transduction mechanisms regulated by the p53 signaling pathway.

## Materials and methods

### Construction of plasmids

Two reporter plasmids, pAAVS1-(*p53RE(C3)*)-Cerulean-NLS-(*SV40)*-mCherry-NLS-2A-Puromycin^R^ and pAAVS1-(*p53RE(CDKN1A)*)-Venus-NLS-Neomycin^R^, were constructed as shown in Fig. S1A,B. The former plasmid carried the artificial *p53RE (C3)*, which consists of three tandem repeats of the p53-binding motif (5’-AGACATGTCCGGACATGTCT-3’) with a 16 bp spacer (5’-ACTAGCGGCTGTCACT-3’). Using this plasmid, we can estimate the activation status of p53 from the expression level of the cyan fluorescent proteinwith three tandem repeats of the nuclear localization signal (PKKKRKV×3) from simian virus large T-antigen in the nucleus (Cerulean-NLS). This plasmid also contains the expression unit of (*SV40*)-mCherry-NLS-2A-Puromycin^R^. The 2A peptide sequence (GSGEGRGSLLTCGDVEENPG / P) derived from the *Thosea asigna virus* interrupts the normal peptide bond and leads to ribosomal skipping (Szymczak et al., 2004). Thus, the red fluorescent protein (mCherry-NLS) and puromycin-resistance gene product can be separately expressed in a p53-independent manner. The latter plasmid carried the *p53RE* derived from the *CDKN1A (p21)* gene, which was originally reported in *Cancer Res*. (GenBank accession no. U24170, nucleotides 2241-3258) (Shimada et al., 1999). Using this plasmid, a yellow fluorescent protein with an NLS sequence (Venus-NLS) is p53-dependently expressed in the nucleus. Both plasmids contain the left and right homology arm to the AAVS1 genomic integration site derived from the AAVS1 SA-2A-puro-pA donor plasmid (Addgene) (Kotin et al., 1992). The mRNA degradation signal, AU-rich element (ATTTATTT ATTTATTT ATTTA), is inserted in the 3’-noncoding region of Venus-NLS and Cerulean-NLS to monitor the quick response in p53-dependent transcriptional activity (Chen and Shyu, 1995).

Five reporter plasmids were also constructed by replacing *p53RE(CDKN1A)* of pAAVS1-(*p53RE(CDKN1A)*)-Venus-NLS-Neomycin^R^ to *p53RE(each gene)*; each gene is *BAX*, *MDM2*, *GADD45*, *RRM2B*, *14-3-3σ*. The following *p53RE* regions were used for each gene, based on the transcription start point: *BAX*; −896 to −332, *MDM2*; 594 to 801, *GADD45*; 1388 to 1721, *RRM2B*; 1953 to 2584, *14-3-3σ*; −2553 to −1768 (Kato et al., 2003; Shimada et al., 1999).

Using the pAAVS1-(*p53RE(CDKN1A)*)-Venus-NLS-Neomycin^R^, we also constructed a cell cycle reporter plasmid, p*AAVS1*-(*p53RE(CDKN1A)*)-Venus-NLS-(*SV40)*-mCherry-NLS-2A-Puromycin^R^-2A-Fucci-G1-Cerulean (Fig. S6A). This plasmid can separately express the mCherry-NLS, puromycin-resistance gene product, and Fucci-G1-Cerulean, which is Cerulean fused to a part of the human Cdt1 of pFucci-G1 Orange (Amalgaam, MBL) (Sakaue-Sawano et al., 2008).

The plasmid pBS-p53Nn-Zeo^R^-p53Cc for making the p53-knockout cell line carried the 974 bp DNA fragment of the 5’-noncoding region in the human p53 gene (*TP53*) from nucleotide number −971 to 4 and the 869 bp DNA fragment of the 3’-noncoding region in *TP53* starting from the stop codon. Using this plasmid, we can replace the p53 coding region with the Zeocin-resistance gene of pcDNA3.1/Zeo (Invitrogen).

The guide RNA expression plasmids for the CRISPR/Cas9 system were constructed using pX330 from Addgene. The pX330-AAVS1-1, -p53N, or -p53C carried the DNA fragment, 5’-GTC CCC TCC ACC CCA CAG TG-3’, 5’-GCG GGT CAC TGC CAT GGA GG-3’, or 5’-GGA GAA TGT CAG TCT GAG TC-3’, between two BbsI sites in pX330, respectively.

### Cell line and cell culture

A549 cells (p53 wild type) were obtained from American type tissue culture collection (ATCC) (Rockville, MD, USA). Cells were cultured on a 35 mm dish in Dulbecco’s modified Eagle’s medium (DMEM)-high glucose (Sigma Aldrich) supplemented with 10% fetal bovine serum (FBS) (Japan Bioserum), 100 units/ml penicillin, and 100 μg/ml streptomycin (Thermo Fisher). For stress addition, cells were treated with 10 μg/ml etoposide (Wako) or irradiated with 25 J/m^2^ UV-C (254 nm) by FUNA UV Crosslinker, FS-800 (Funakoshi). All cells were maintained in a humidified atmosphere of 5% CO2 at 37°C.

### Establishment of knock-in and knockout cell lines

For the establishment of the reporter cell line, A549 cells were plated on a 35 mm dish for 24 h and transfected with 0.2 μg of pX330-AAVS1-1 vector and 0.4 μg knock-in donors by using Lipofectamine 2000 (Invitrogen). At 24 h after transfection, the cells were seeded onto a 35 mm dish and treated with 0.5 μg/ml puromycin (Invivogen)/1 mg/ml G418 (Roche) for more than 14 days. The culture medium was changed every 3 ~ 5 days. The selected cell lines were subcloned with a 96-well plate. For the establishment of the p53-knockout cell line, A549[AAVS1](*p53RE(C3)*)-Cerulean-NLS-(*SV40*)-mCherry-NLS / (*p53RE(CDKN1A)*)-Venus-NLS cells were plated on a 35 mm dish for 24 h and transfected with 0.6 μg of pBS-p53Nn-Zeo^R^-p53Cc, 0.5 μg of pX330-p53N, and 0.5 μg of pX330–p53C using 4 μl of Lipofectamine 2000, selected with 100 μg/ ml Zeocin (Invivogen), and subcloned with a 96-well plate.

### Cell cycle synchronization

A double thymidine block was used for S phase synchronization and serum starvation for G1 phase synchronization (Bostock et al., 1971; Johmura et al., 2014). For the double thymidine block, A549 reporter cells were treated with 2 mM thymidine for 12 h, released for 12 h, and then treated with 2 mM thymidine for 12 h again before stress addition. For serum starvation, A549 cells were treated with DMEM without FBS for 48 h before stress addition.

### Live-cell imaging

Live-cell imaging was performed with BIOREVO BZ-X710 (Keyence) for the data in Fig. 3, 4 and BIOREVO BZ-9000 (Keyence) for other data. The following filter sets were used for observation: for mCherry, TexasRed (excitation filter: 560/40 nm; emission filter: 630/60 nm; dichroic mirror: 595 nm, for BIOREVO BZ-X710) or TRITC (excitation filter: 540/25 nm; emission filter: 605/55 nm; dichroic mirror: 565 nm, for BIOREVO BZ-9000); for Venus, YFP (excitation filter: 500/24 nm; emission filter: 542/27 nm; dichroic mirror: 520 nm); and for Cerulean, CFP (excitation filter: 448/20 nm; emission filter: 482/25 nm; dichroic mirror: 458 nm). All live-cell imaging was performed with a Plan Fluor D 10x / NA030 objective lens (Nikon).

### Image analysis

The transcriptional activity from different promoters was monitored by three fluorescent proteins fused with NLS, so that the transcriptional activity was detected as the fluorescence intensity present in the nucleus. Quantitative analysis of fluorescence intensity were performed as previously described (Imagawa et al., 2009). The detailed procedure used for detection and quantification of fluorescent signals with Image Analysis was as follows: 1) The original image is smoothed by applying a Gaussian filter, and then the background signal is subtracted; 2) Based on the fluorescence image of mCherry, areas with values above a set threshold are detected as nuclei and 3) The fluorescence intensity of each reporter is calculated based on the detected nuclear area. Single cells were tracked manually by the phase and fluorescence images. Cell death was determined manually from the changes of cell morphology by phase images (Fig. S3A). Only experimental data under UV-C irradiation conditions (Fig. 4) were analyzed using Fiji and LIMTracker (Aragaki et al., 2022; Schindelin et al., 2012).

### Immunofluorescence

A549 cells were plated on a 35 mm dish and their cell cycles were synchronized (the detailed schedules are shown in Fig. 5A), and then etoposide stimulation was performed for 24 h. Cells were fixed with 10% neutral buffered formalin solution (Wako) for 15 min, washed in phosphate-buffered saline (PBS), permeabilized with 0.2% Triton X-100 / PBS for 15 min, and blocked with 10% FBS/PBS for 1 h. After that, cells were incubated with the first antibody/1% FBS/PBS for 16 h at 4°C and incubated with the 2nd antibody/DAPI (Wako)/1% FBS/PBS for 30 min at room temperature. The first antibody was mouse monoclonal anti-p53/DO-1 (SC-126) (Santa Cruz Biotechnology) for determination of the p53 expression level and mouse monoclonal anti-p53(Ser15)/16G8 (9286) (Cell Signaling) for determination of the p53 Ser15 phosphorylation level. The second antibody was AlexaFluoro488 anti-mouse IgG (A-11001) (Invitrogen). Cells were imaged on a BIOREVO BZ-9000 fluorescent microscope (Keyence). The following filter sets were used for observation: for AlexaFluoro488, GFP (excitation filter: 470/40 nm; emission filter: 535/50 nm; dichroic mirror: 495 nm); and for DAPI, DAPI (excitation filter: 360/40 nm; emission filter: 460/50 nm; dichroic mirror: 400 nm). Nuclei were detected by DAPI images and the relative expression levels of proteins in the nuclear region were quantified.

### Cell cycle classification

Cell traces were classified automatically by the cell cycle phase at drug treatment based on the expression pattern of Fucci-G1-Cerulean and the time of cell division (Fig. 6C). For cells classified to G1 phase, those that were treated within the first 6 h after cell division and those that were treated after the first 6 h following cell division were further classified as early G1 and late G1 cells, respectively. For cells classified to S/G2 phase, those that were treated within the first 6 h after the peak and those that were treated after the first 6 h following the peak were further classified as S and G2 cells, respectively. The time window of each cell cycle phase was decided by the measured parameters of cell cycle length and cell cycle population of A549 cells (Fig. S6B,C) (cell cycle length: 21.9±0.2 h; cell cycle population: G1 (51.4±2.4%), S (35.2±1.7%), G2 (12.3±0.3%)). Specifically, we calculated the length of the S phase and obtained the value of 7.4 h, and therefore we separated cells that were treated first 6 h after the peak or not were classified S or G2 cells, respectively.

## Acknowledgments

This work was supported in part through Hokkaido University, Global Facility Center (GFC), Pharma Science Open Unit (PSOU), found by MEXT under “Support Program for Implementation of New Equipment Sharing System”.

## Competing interests

The authors declare no competing interests.

## Funding

This work was supported in part by a Grant-in-Aid for Scientific Research(B) (20H02873 to K.S.) and the Photo-excitonix Project at Hokkaido University (to K.S.).

## Data availability

All relevant data are within the manuscript and its Supplementary Information files.

## Supplementary information

This article contains supplementary information.

## List of Symbols and Abbreviations

The abbreviations used are:

*p53RE*: p53 responsive element
*TP53*: tumor protein 53
ERK: extracellular signal-regulated kinase
NF-κB: nuclear factor-kappa B
NFAT: Nuclear factor of activated T-cells
*AAVS1*: Adeno-associated virus integration site 1
NLS: nuclear localization signal
*SV40*: simian virus 40 promoter
Puromycin^R^: puromycin resistance
*CDKN1A*: cyclin dependent kinase inhibitor 1A
Neomycin^R^: neomycin resistance
*BAX*: Bcl-2-associated X protein
*MDM2*: murine double minute 2
*GADD45*: growth arrest and DNA-damage-inducible protein
*RRM2B*: ribonucleoside-diphosphate reductase subunit M2 B
UV-C: ultraviolet-C
JNK: c-Jun N-terminal kinase
Fucci: fluorescent ubiquitination-based cell cycle indicator
Cdt1: chromatin licensing and DNA replication factor 1
Zeo^R^: zeocin resistance
CRISPR: clustered regularly interspaced short palindromic repeats
DMEM: Dulbecco’s modified Eagle’s medium
FBS: fetal bovine serum
TRITC: tetramethylrhodamine
YFP: yellow fluorescent protein
CFP: cyan fluorescent protein
DAPI: 4’,6-diamidino-2-phenylindole
DSB: double-strand break
DTB: double thymidine block
SS: serum starvation
CDK1: cyclin-dependent kinase 1
p53S15P: p53 phosphorylated at Ser15
ARC: apoptosis repressor with the caspase recruitment domain

## Notes

### Competing Interest Statement

The authors have declared no competing interest.

### Summary of Updates

We added new results for UV-C treated conditions (Newly added as Figure 4); We also revised the main text; Supplemental files updated.

## References

Appella, E. and Anderson, C. W. (2001). Post-translational modifications and activation of p53 by genotoxic stresses. Eur. J. Biochem. 268, 2764–2772.

Aragaki, H., Ogoh, K., Kondo, Y. and Aoki, K. (2022). LIM Tracker: a software package for cell tracking and analysis with advanced interactivity. Sci. Rep. 12, 2702.

Bieging, K. T. and Attardi, L. D. (2012). Deconstructing p53 transcriptional networks in tumor suppression. Trends Cell Biol. 22, 97–106.

Bornstein, G., Bloom, J., Sitry-Shevah, D., Nakayama, K., Pagano, M. and Hershko, A. (2003). Role of the SCFSkp2 Ubiquitin Ligase in the Degradation of p21Cip1 in S Phase*. J. Biol. Chem. 278, 25752–25757.

Bostock, C. J., Prescott, D. M. and Kirkpatrick, J. B. (1971). An evaluation of the double thymidine block for synchronizing mammalian cells at the G1-S border. Experimental Cell Research 68, 163–168.

Buschmann, T., Potapova, O., Bar-Shira, A., Ivanov, V. N., Fuchs, S. Y., Henderson, S., Fried, V. A., Minamoto, T., Alarcon-Vargas, D., Pincus, M. R., et al. (2001). Jun NH2-terminal kinase phosphorylation of p53 on Thr-81 is important for p53 stabilization and transcriptional activities in response to stress. Mol. Cell. Biol. 21, 2743–2754.

Chen, C. Y. and Shyu, A. B. (1995). AU-rich elements: characterization and importance in mRNA degradation. Trends Biochem. Sci. 20, 465–470.

Chen, X., Chen, J., Gan, S., Guan, H., Zhou, Y., Ouyang, Q. and Shi, J. (2013). DNA damage strength modulates a bimodal switch of p53 dynamics for cell-fate control. BMC Biol. 11, 73.

Fischer, M. (2017). Census and evaluation of p53 target genes. Oncogene 36, 3943–3956.

Foo, R. S.-Y., Nam, Y.-J., Ostreicher, M. J., Metzl, M. D., Whelan, R. S., Peng, C.-F., Ashton, A. W., Fu, W., Mani, K., Chin, S.-F., et al. (2007). Regulation of p53 tetramerization and nuclear export by ARC. Proc. Natl. Acad. Sci. U. S. A. 104, 20826–20831.

Gaglia, G. and Lahav, G. (2014). Constant rate of p53 tetramerization in response to DNA damage controls the p53 response. Mol. Syst. Biol. 10, 753.

Gaglia, G., Guan, Y., Shah, J. V. and Lahav, G. (2013). Activation and control of p53 tetramerization in individual living cells. Proc. Natl. Acad. Sci. U. S. A. 110, 15497–15501.

Gartel, A. L. and Radhakrishnan, S. K. (2005). Lost in transcription: p21 repression, mechanisms, and consequences. Cancer Res. 65, 3980–3985.

Goswami, P. C., Roti Roti, J. L. and Hunt, C. R. (1996). The cell cycle-coupled expression of topoisomerase IIalpha during S phase is regulated by mRNA stability and is disrupted by heat shock or ionizing radiation. Mol. Cell. Biol. 16, 1500–1508.

Granada, A. E., Jiménez, A., Stewart-Ornstein, J., Blüthgen, N., Reber, S., Jambhekar, A. and Lahav, G. (2020). The effects of proliferation status and cell cycle phase on the responses of single cells to chemotherapy. Mol. Biol. Cell 31, 845–857.

Hainsworth, J. D. and Greco, F. A. (1995). Etoposide: twenty years later. Ann. Oncol. 6, 325–341.

Haneke, K., Schott, J., Lindner, D., Hollensen, A. K., Damgaard, C. K., Mongis, C., Knop, M., Palm, W., Ruggieri, A. and Stoecklin, G. (2020). CDK1 couples proliferation with protein synthesis. J. Cell Biol. 219,.

Harton, M. D., Koh, W. S., Bunker, A. D., Singh, A. and Batchelor, E. (2019). p53 pulse modulation differentially regulates target gene promoters to regulate cell fate decisions. Mol. Syst. Biol. 15, e8685.

Horn, H. F. and Vousden, K. H. (2007). Coping with stress: multiple ways to activate p53. Oncogene 26, 1306–1316.

Hsiang, Y. H., Wu, H. Y. and Liu, L. F. (1988). Proliferation-dependent regulation of DNA topoisomerase II in cultured human cells. Cancer Res. 48, 3230–3235.

Imagawa, T., Terai, T., Yamada, Y., Kamada, R. and Sakaguchi, K. (2009). Evaluation of transcriptional activity of p53 in individual living mammalian cells. Anal. Biochem. 387, 249–256.

Johmura, Y., Shimada, M., Misaki, T., Naiki-Ito, A., Miyoshi, H., Motoyama, N., Ohtani, N., Hara, E., Nakamura, M., Morita, A., et al. (2014). Necessary and sufficient role for a mitosis skip in senescence induction. Mol. Cell 55, 73–84.

Kamada, R., Toguchi, Y., Nomura, T., Imagawa, T. and Sakaguchi, K. (2016). Tetramer formation of tumor suppressor protein p53: Structure, function, and applications. Biopolymers 106, 598–612.

Karni-Schmidt, O., Lokshin, M. and Prives, C. (2016). The Roles of MDM2 and MDMX in Cancer. Annu. Rev. Pathol. 11, 617–644.

Kastenhuber, E. R. and Lowe, S. W. (2017). Putting p53 in Context. Cell 170, 1062–1078.

Kato, S., Han, S.-Y., Liu, W., Otsuka, K., Shibata, H., Kanamaru, R. and Ishioka, C. (2003). Understanding the function-structure and function-mutation relationships of p53 tumor suppressor protein by high-resolution missense mutation analysis. Proc. Natl. Acad. Sci. U. S. A. 100, 8424–8429.

Kotin, R. M., Linden, R. M. and Berns, K. I. (1992). Characterization of a preferred site on human chromosome 19q for integration of adeno-associated virus DNA by non-homologous recombination. EMBO J. 11, 5071–5078.

Krenning, L., Feringa, F. M., Shaltiel, I. A., van den Berg, J. and Medema, R. H. (2014). Transient activation of p53 in G2 phase is sufficient to induce senescence. Mol. Cell 55, 59–72.

Lee, T. K., Denny, E. M., Sanghvi, J. C., Gaston, J. E., Maynard, N. D., Hughey, J. J. and Covert, M. W. (2009). A noisy paracrine signal determines the cellular NF-kappaB response to lipopolysaccharide. Sci. Signal. 2, ra65.

Levrero, M., De Laurenzi, V., Costanzo, A., Gong, J., Wang, J. Y. and Melino, G. (2000). The p53/p63/p73 family of transcription factors: overlapping and distinct functions. J. Cell Sci. 113 (Pt 10), 1661–1670.

Li, Y.-Z., Lu, D.-Y., Tan, W.-Q., Wang, J.-X. and Li, P.-F. (2008). p53 initiates apoptosis by transcriptionally targeting the antiapoptotic protein ARC. Mol. Cell. Biol. 28, 564–574.

Loewer, A., Batchelor, E., Gaglia, G. and Lahav, G. (2010). Basal dynamics of p53 reveal transcriptionally attenuated pulses in cycling cells. Cell 142, 89–100.

Loffreda, A., Jacchetti, E., Antunes, S., Rainone, P., Daniele, T., Morisaki, T., Bianchi, M. E., Tacchetti, C. and Mazza, D. (2017). Live-cell p53 single-molecule binding is modulated by C-terminal acetylation and correlates with transcriptional activity. Nat. Commun. 8, 313.

Marshall, C. J. (1995). Specificity of receptor tyrosine kinase signaling: transient versus sustained extracellular signal-regulated kinase activation. Cell 80, 179–185.

Meek, D. W. (2009). Tumour suppression by p53: a role for the DNA damage response? Nat. Rev. Cancer 9, 714–723.

Miura, H., Kondo, Y., Matsuda, M. and Aoki, K. (2018). Cell-to-Cell Heterogeneity in p38-Mediated Cross-Inhibition of JNK Causes Stochastic Cell Death. Cell Rep. 24, 2658–2668.

Murray-Zmijewski, F., Slee, E. A. and Lu, X. (2008). A complex barcode underlies the heterogeneous response of p53 to stress. Nat. Rev. Mol. Cell Biol. 9, 702–712.

Muslimović, A., Nyström, S., Gao, Y. and Hammarsten, O. (2009). Numerical analysis of etoposide induced DNA breaks. PLoS One 4, e5859.

Nagai, T., Ibata, K., Park, E. S., Kubota, M., Mikoshiba, K. and Miyawaki, A. (2002). A variant of yellow fluorescent protein with fast and efficient maturation for cell-biological applications. Nat. Biotechnol. 20, 87–90.

Nelson, D. E., Ihekwaba, A. E. C., Elliott, M., Johnson, J. R., Gibney, C. A., Foreman, B. E., Nelson, G., See, V., Horton, C. A., Spiller, D. G., et al. (2004). Oscillations in NF-kappaB signaling control the dynamics of gene expression. Science 306, 704–708.

Nitiss, J. L. (2009a). Targeting DNA topoisomerase II in cancer chemotherapy. Nat. Rev. Cancer 9, 338–350.

Nitiss, J. L. (2009b). DNA topoisomerase II and its growing repertoire of biological functions. Nat. Rev. Cancer 9, 327–337.

Oleinik, N. V., Krupenko, N. I. and Krupenko, S. A. (2007). Cooperation between JNK1 and JNK2 in activation of p53 apoptotic pathway. Oncogene 26, 7222–7230.

Oliner, J. D., Pietenpol, J. A., Thiagalingam, S., Gyuris, J., Kinzler, K. W. and Vogelstein, B. (1993). Oncoprotein MDM2 conceals the activation domain of tumour suppressor p53. Nature 362, 857–860.

Olivier, M., Hollstein, M. and Hainaut, P. (2010). TP53 mutations in human cancers: origins, consequences, and clinical use. Cold Spring Harb. Perspect. Biol. 2, a001008.

Paek, A. L., Liu, J. C., Loewer, A., Forrester, W. C. and Lahav, G. (2016). Cell-to-Cell Variation in p53 Dynamics Leads to Fractional Killing. Cell 165, 631–642.

Purvis, J. E. and Lahav, G. (2013). Encoding and decoding cellular information through signaling dynamics. Cell 152, 945–956.

Purvis, J. E., Karhohs, K. W., Mock, C., Batchelor, E., Loewer, A. and Lahav, G. (2012). p53 dynamics control cell fate. Science 336, 1440–1444.

Rizzo, M. A., Springer, G. H., Granada, B. and Piston, D. W. (2004). An improved cyan fluorescent protein variant useful for FRET. Nat. Biotechnol. 22, 445–449.

Roos, W. P., Thomas, A. D. and Kaina, B. (2016). DNA damage and the balance between survival and death in cancer biology. Nat. Rev. Cancer 16, 20–33.

Sadelain, M., Papapetrou, E. P. and Bushman, F. D. (2011). Safe harbours for the integration of new DNA in the human genome. Nat. Rev. Cancer 12, 51–58.

Saha, M. N., Jiang, H., Yang, Y., Zhu, X., Wang, X., Schimmer, A. D., Qiu, L. and Chang, H. (2012). Targeting p53 via JNK pathway: a novel role of RITA for apoptotic signaling in multiple myeloma. PLoS One 7, e30215.

Sakaue-Sawano, A., Kurokawa, H., Morimura, T., Hanyu, A., Hama, H., Osawa, H., Kashiwagi, S., Fukami, K., Miyata, T., Miyoshi, H., et al. (2008). Visualizing spatiotemporal dynamics of multicellular cell-cycle progression. Cell 132, 487–498.

Schindelin, J., Arganda-Carreras, I., Frise, E., Kaynig, V., Longair, M., Pietzsch, T., Preibisch, S., Rueden, C., Saalfeld, S., Schmid, B., et al. (2012). Fiji: an open-source platform for biological-image analysis. Nat. Methods 9, 676–682.

Shaner, N. C., Campbell, R. E., Steinbach, P. A., Giepmans, B. N. G., Palmer, A. E. and Tsien, R. Y. (2004). Improved monomeric red, orange and yellow fluorescent proteins derived from Discosoma sp. red fluorescent protein. Nat. Biotechnol. 22, 1567–1572.

Shimada, A., Kato, S., Enjo, K., Osada, M., Ikawa, Y., Kohno, K., Obinata, M., Kanamaru, R., Ikawa, S. and Ishioka, C. (1999). The Transcriptional Activities of p53 and Its Homologue p51/p63: Similarities and Differences. Cancer Res. 59, 2781–2786.

Stewart-Ornstein, J. and Lahav, G. (2016). Dynamics of CDKN1A in Single Cells Defined by an Endogenous Fluorescent Tagging Toolkit. Cell Rep. 14, 1800–1811.

Szymczak, A. L., Workman, C. J., Wang, Y., Vignali, K. M., Dilioglou, S., Vanin, E. F. and Vignali, D. A. A. (2004). Correction of multi-gene deficiency in vivo using a single “self-cleaving” 2A peptide-based retroviral vector. Nat. Biotechnol. 22, 589–594.

Tammaro, M., Barr, P., Ricci, B. and Yan, H. (2013). Replication-dependent and transcription-dependent mechanisms of DNA double-strand break induction by the topoisomerase 2-targeting drug etoposide. PLoS One 8, e79202.

Weinberg, R. L., Veprintsev, D. B., Bycroft, M. and Fersht, A. R. (2005). Comparative binding of p53 to its promoter and DNA recognition elements. J. Mol. Biol. 348, 589–596.

Wu, M., Ye, H., Tang, Z., Shao, C., Lu, G., Chen, B., Yang, Y., Wang, G. and Hao, H. (2017). p53 dynamics orchestrates with binding affinity to target genes for cell fate decision. Cell Death Dis. 8, e3130.

Yang, R., Huang, B., Zhu, Y., Li, Y., Liu, F. and Shi, J. (2018). Cell type-dependent bimodal p53 activation engenders a dynamic mechanism of chemoresistance. Sci Adv 4, eaat5077.

Yissachar, N., Sharar Fischler, T., Cohen, A. A., Reich-Zeliger, S., Russ, D., Shifrut, E., Porat, Z. and Friedman, N. (2013). Dynamic response diversity of NFAT isoforms in individual living cells. Mol. Cell 49, 322–330.

