## Supplementary Information for "Development of a fluorescence reporter system to quantify transcriptional activity of endogenous p53 in living cells"

**Contents**

**Materials and methods**

**Figure S1: Design and characterization of the reporter system.**

**Figure S2: Correlation of transcriptional activation of different target genes and internal standard of p53 activation status.**

**Figure S3: Correlation of p53-independent transcriptional activity and p53-dependent transcriptional activity or cell death.**

**Figure S4: Single cell traces of all cells classified by the time of cell division.**

**Figure S5: Temporal changes of the growth rate and transcriptional activity of endogenous p53 by cell cycle synchronization.**

**Figure S6: Development of the reporter system for monitoring the transcriptional activity of p53 and the cell cycle.**

### Materials and methods

Western blotting
A549 reporter cells (p53 wild type and p53-knockout) were plated in a 35 mm dish for 24 h, and then etoposide stimulation was performed. Cells were harvested immediately at 0, 12, 24, or 48 h after stimulation. Total cell extracts were sampled with 1× sample buffer (50 mM Tris-HCl, pH 6.8, 10% glycerol, 2% SDS, 6% 2-mercaptoethanol) and cell lysates were normalized for total protein. After sonication, samples were separated by SDS-PAGE and transferred to polyvinylidene difluoride membranes. Proteins were detected by enhanced chemiluminescence with the antibodies. The first antibody was mouse monoclonal anti-p53/DO-1 (SC-126) (Santa Cruz Biotechnology) and the second antibody was an anti-mouse IgG-HRP-linked antibody (NA931) (GE Healthcare).

Genome PCR
A549 reporter cells (p53 wild type and p53-knockout) were plated in a 35 mm dish for 24 h, and genome DNA was harvested by DNAzol (Cosmo Bio). Primer sequences were as follows: for *TP53* N-terminal verification, 5'-AGTCTGCACGGGAAGGAGCCTACCCCCATG-3' and 5'-AAACGAACGTTGTTTTCAGGAAGT-3'; for *TP53* C-terminal verification, 5'-CGCCATAAAAAACTCATGTTCAAG-3' and 5'-GGCTGGCCATGGTGGC ATGAACCTGTGGTC-3'.

Growth rate measurement
A549 reporter cells were plated on a 35 mm dish for 24 h and cell cycle synchronization was performed as described in the *Experimental procedures*. Cells in the same area were imaged every 6 h (SS) or 3 h (DTB) and nuclei were detected by mCherry fluorescence. The growth rate was calculated by the number of detected nuclei.

Doubling-time measurement
A549 reporter cells were plated on a 35 mm dish for 24 h and then cells in the same area were imaged every 6 h. The growth rate was calculated by the number of detected nuclei (see Supporting Information Fig. S5 for detail).

Flow cytometric analysis
A549 cells were plated in a 10 cm dish for 2 days. Cells were harvested, washed with 2% FBS/PBS, and fixed with ice-cold 99.5% ethanol. The fixed cells were washed with 2% FBS/PBS and labeled by using an Annexin-V-FLUOS Staining Kit (Roche) for 15 min, followed by analysis with a flow cytometer (Gallios, Beckman Coulter, Miami, FL, USA). Data were analyzed by FlowJo software.

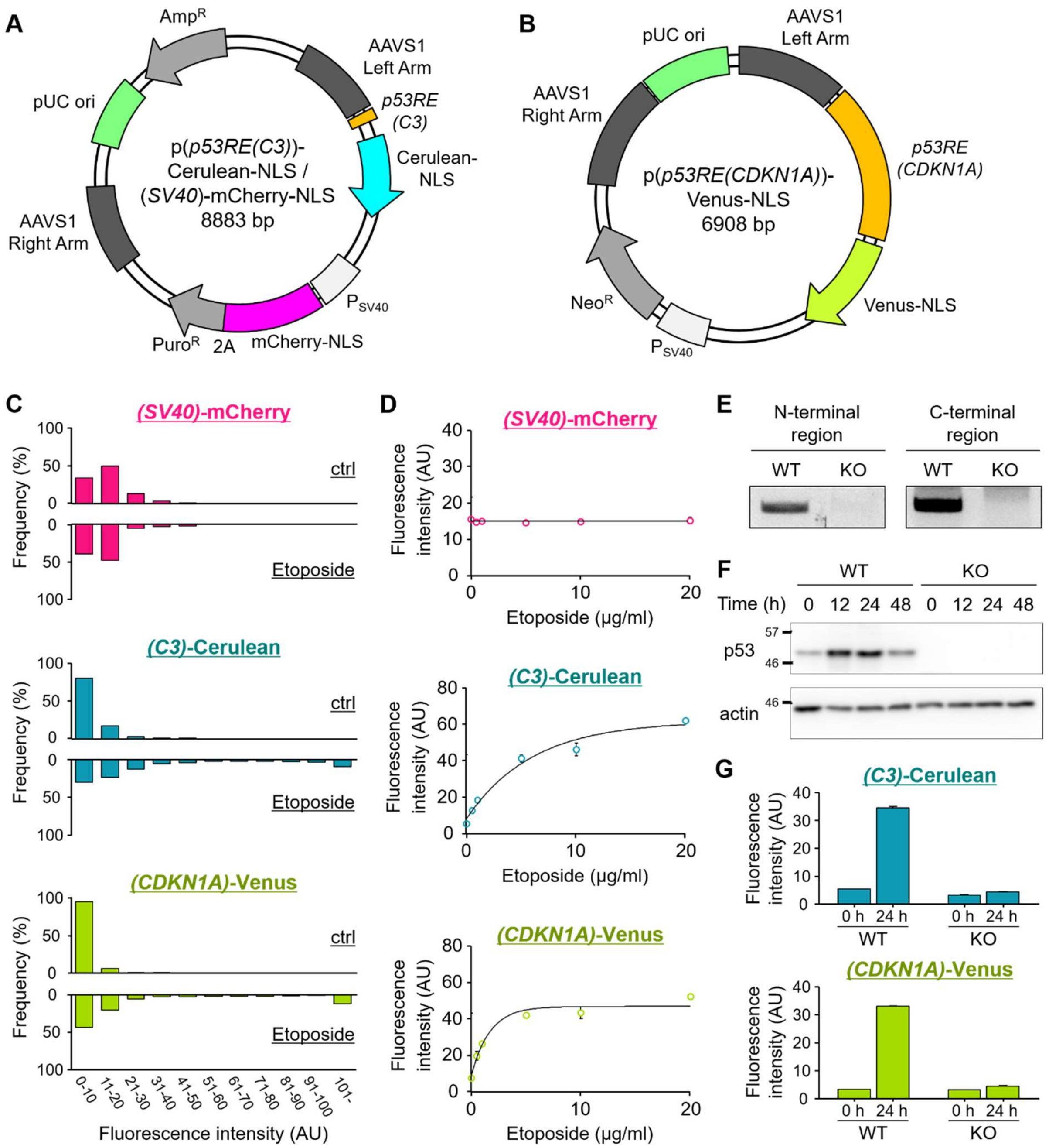


**Figure S1 Design and characterization of the reporter system.**

A) The reporter vector, p(*p53RE(C3)*)-Cerulean-NLS, (*SV40*)-mCherry-NLS, for quantifying the transcriptional activity of the endogenous p53 to the artificial p53 response element.

B) The reporter vector, p(*p53RE(CDKN1A)*)-Venus-NLS, for quantifying the transcriptional activity of the endogenous p53 to the p53 response element (*p53RE*). The sequence of *p53RE* was derived from the *CDKN1A* gene (-2,324~ -1,311 from the TSS site; 1,014 bp).

C) Distribution of fluorescence intensity at 24 h with (+ Etp) or without (ctrl) treatment with 10 μg/ml etoposide.

D) Dependence of the reporter cell line on the etoposide concentration. The plots show the mean fluorescence intensity and S.D. at 24 h after etoposide treatment from three independent experiments (more than 400 cells were analyzed in each experiment).

E) Genome PCR of the p53 genomic region. WT and KO indicate the p53 wild-type reporter cell line and p53-knockout reporter cell line, respectively.

F) Time course of the p53 expression level after 10 μg/ml etoposide treatment by western blotting.

G) Fluorescence intensity at 0 h and 24 h after 10 μg/ml etoposide treatment. Horizontal lines indicate the mean with S.D. More than 4,000 cells were analyzed under each condition.

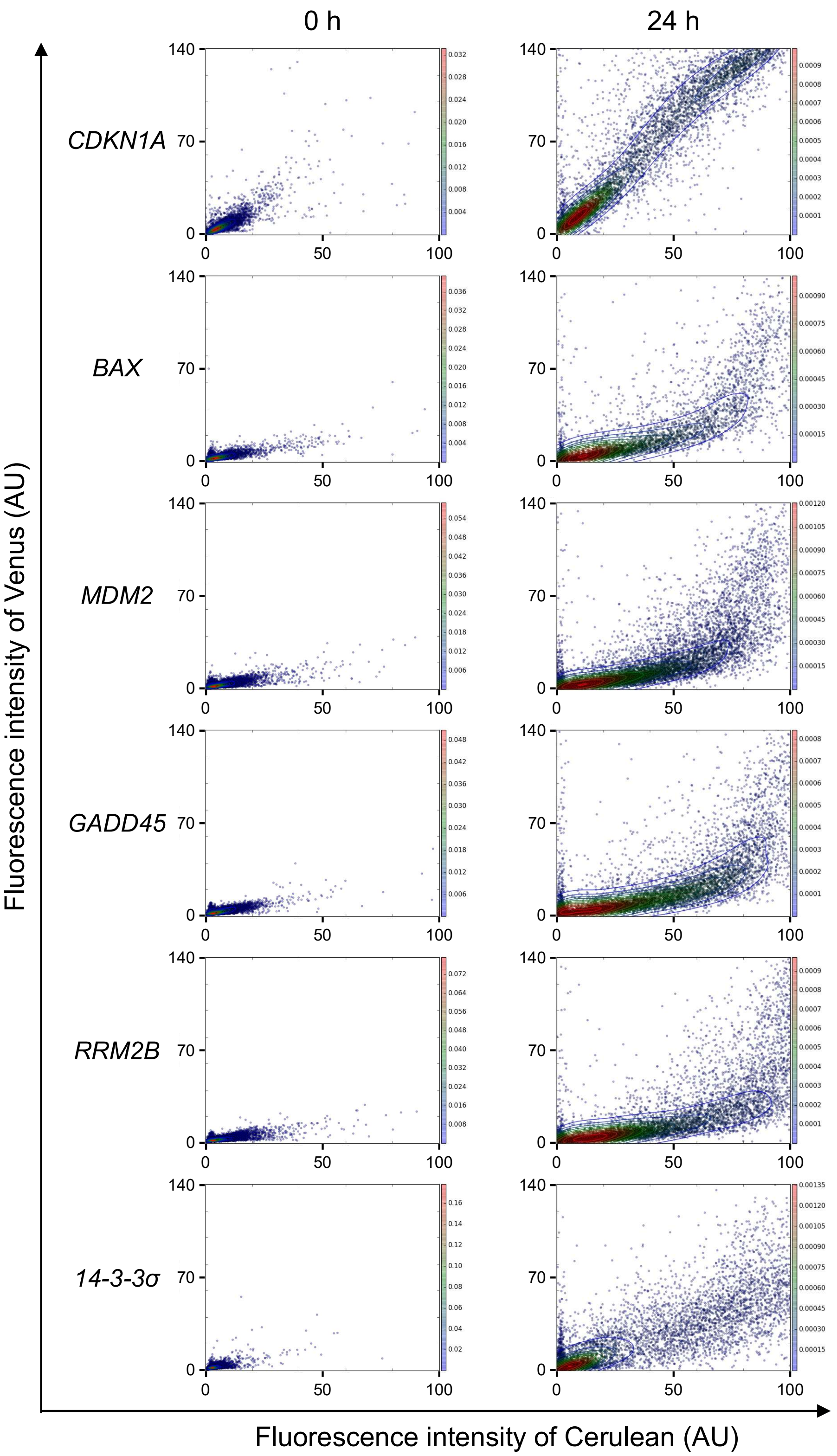


**Figure S2 Correlation of transcriptional activation of different target genes and internal standard of p53 activation status.**

Correlation analysis of the transcriptional activity for six target genes (Venus) vs internal standard of p53 activation status (Cerulean) at 0 h (left panel) and 24 h (right panel) after 10 μg/ml etoposide treatment. The target genes corresponding to the *p53RE*s used in the reporter cell lines are *CDKN1A*, *BAX*, *MDM2*, *GADD45*, *RRM2B*, and *14-3-3σ* from top to bottom. Contour lines were calculated and drawn using kernel density estimation.

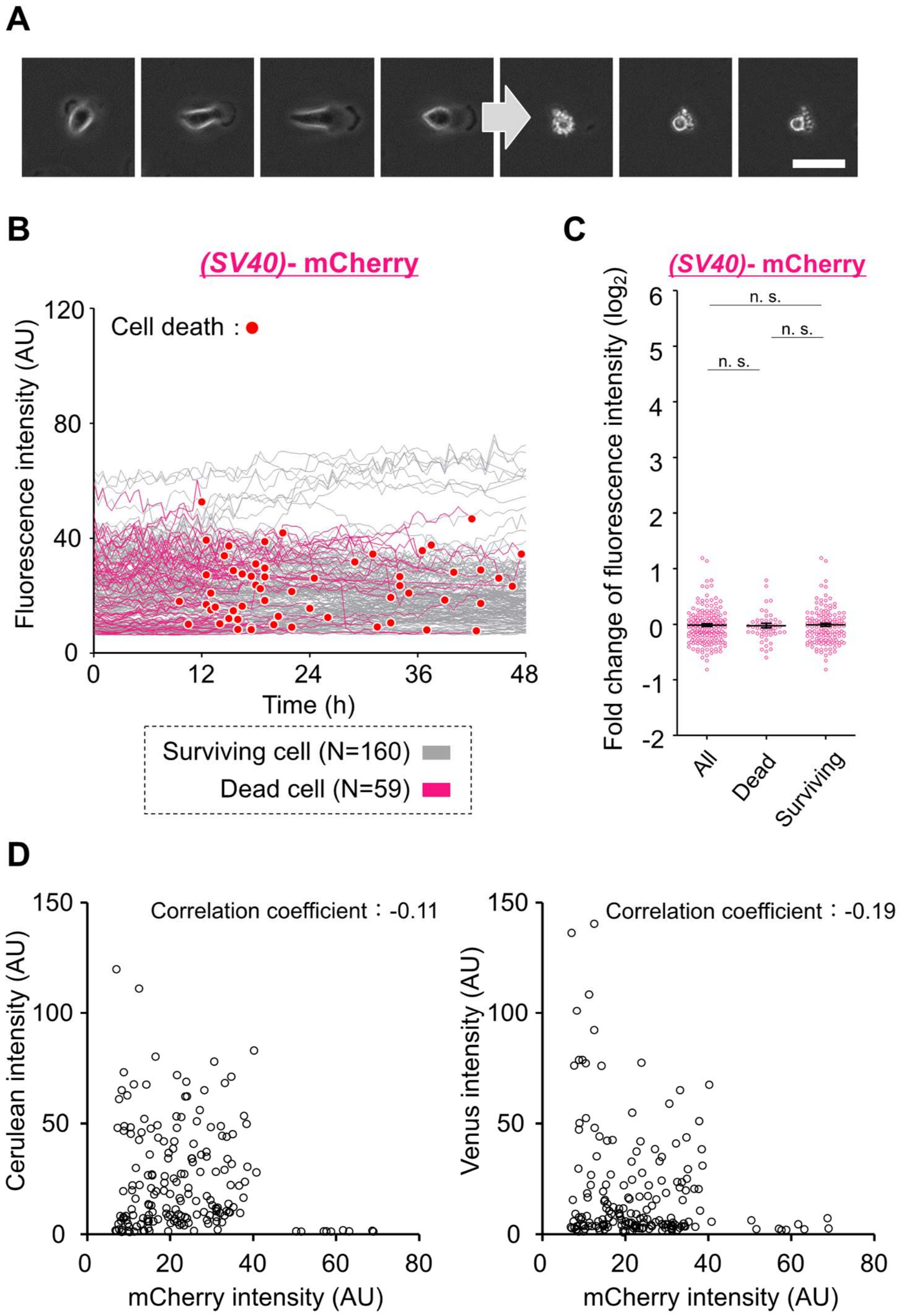


**Figure S3 Correlation of p53-independent transcriptional activity and p53-dependent transcriptional activity or cell death.**

A) Morphological changes of cell death. Cell images were obtained every 30 min.

B) Single cell traces (mCherry) of all cells. Red dots indicate cell death. The gray or colored lines show the traces of surviving cells (n=160) and dying cells (n=59), respectively.

C) Fold changes of fluorescence intensity (mCherry) at 12 h compared with that at 0 h after 10 μg/ml etoposide treatment. Horizontal lines indicate the mean with S.E.M. P values were calculated by an unpaired t-test with Welch's correction. n.s.=not significant ($p > 0.05$).

D) Correlation analysis of the p53 dependent transcriptional activity (Cerulean; left panel or Venus; right panel) vs p53 independent activity (mCherry) at 24 h.

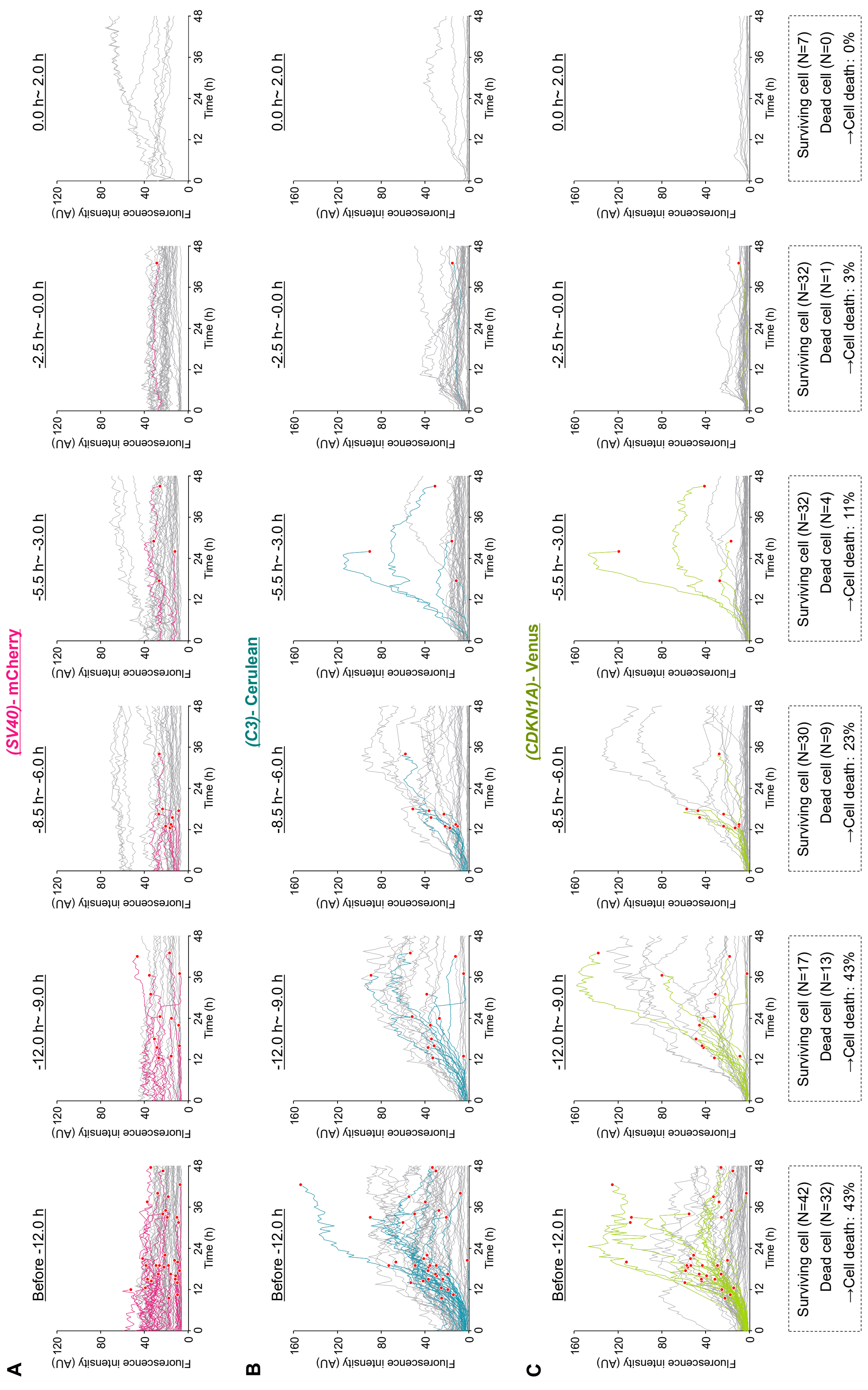


**Figure S4 Single cell traces of all cells classified by the time of cell division.**

A)-C) Single cell traces of the mCherry (A), Cerulean (B), and Venus (C) fluorescence intensity. Red dots indicate cell death. Surviving cells and dying cells are shown by colored lines and gray lines, respectively.

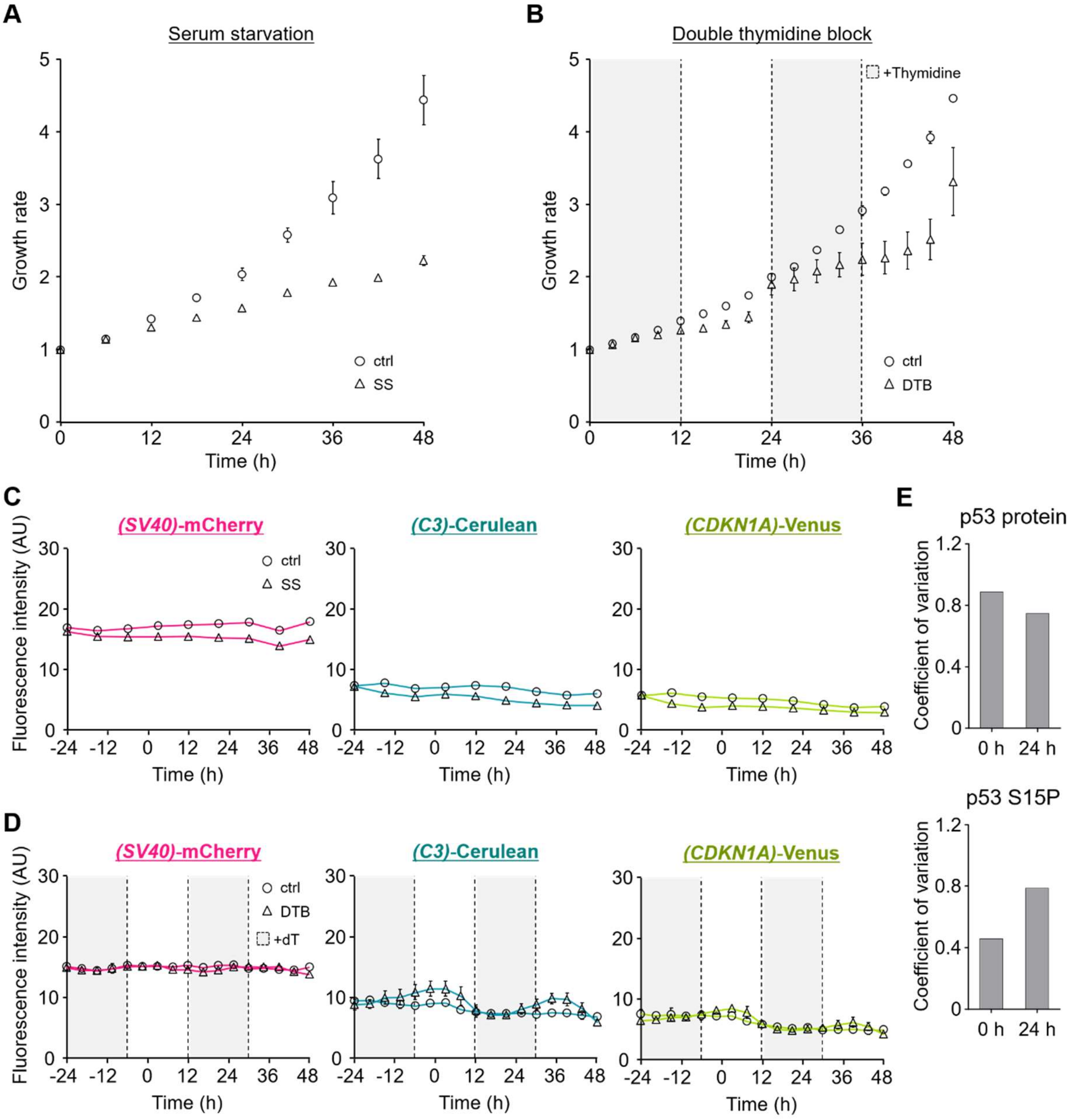


**Figure S5 Temporal changes of the growth rate and transcriptional activity of endogenous p53 by cell cycle synchronization.**

A) The growth rate of the A549 cell line during serum starvation (SS). The plot shows the mean growth rate with S.D. from three independent experiments (more than 1,000 cells were analyzed in each experiment).

B) The growth rate of the A549 cell line during double thymidine block (DTB). The concentration of thymidine was 2 mM. The plot shows the mean growth rate with S.D. from three independent experiments (more than 400 cells were analyzed in each experiment).

C) Temporal changes of the fluorescence intensity during serum starvation (SS). The plot shows the mean fluorescence intensity with S.D. from three independent experiments (more than 1,000 cells were analyzed for each experiment).

D) Temporal changes of the fluorescence intensity during DTB. The concentration of thymidine (dT) was 2 mM. The plot shows the mean fluorescence intensity with S.D. from three independent experiments (more than 400 cells were analyzed in each experiment).

E) Coefficient of variation of p53 expression levels and p53S15P levels at 0 h and 24 h calculated from the data under cell cycle-unsynchronized conditions.

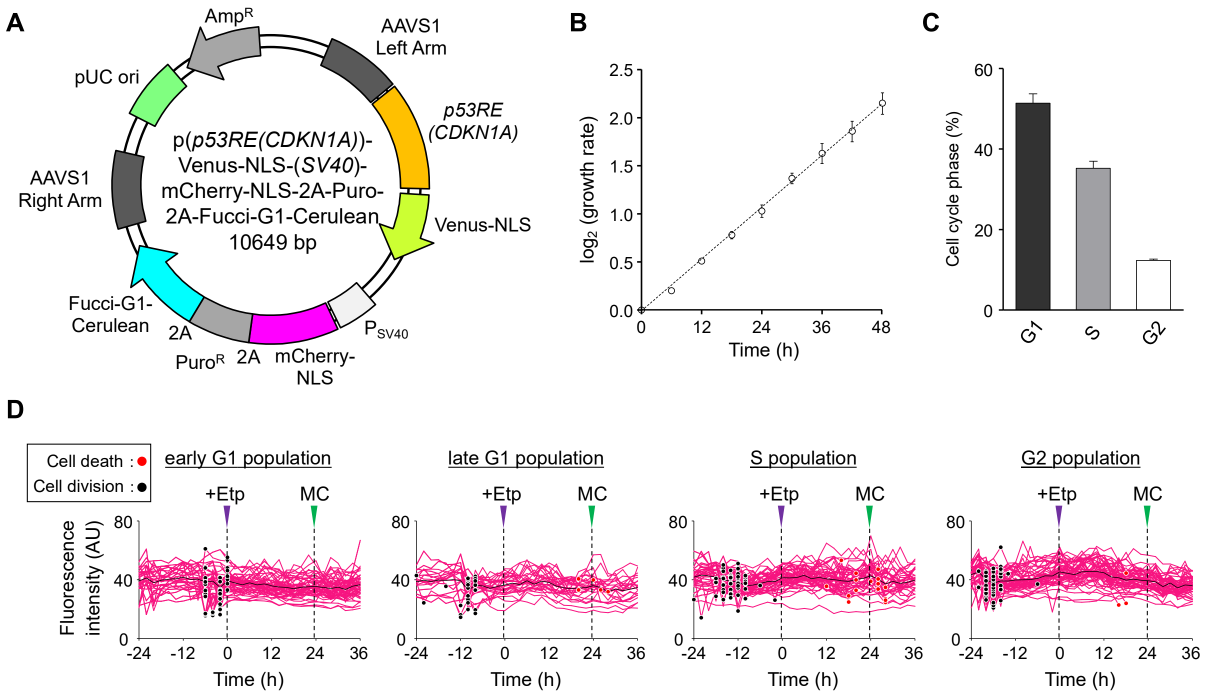


**Figure S8 Development of the reporter system for monitoring the transcriptional activity of p53 and the cell cycle.**

A) The reporter vector, p(*p53RE(CDKN1A)*)-Venus-NLS-(*SV40*)-mCherry-NLS-2A-Puro-2A-Fucci-G1-Cerulean, for quantifying the transcriptional activity of the endogenous p53 to the p53 response element (*p53RE*) and cell cycle. The sequence of *p53RE* is the same as that of the reporter plasmid shown in Supporting Information Fig. S1B.

B) Doubling-time measurement. The plot shows the mean growth rate with S.D. from three independent experiments (more than 1,000 cells were analyzed in each experiment).

C) The ratio of the cell cycle phase quantified by flow cytometry.

D) Single cell traces (mCherry) of all cells classified by the cell cycle phase at stress addition. Black and red dots indicate cell division and cell death, respectively. The black line is a trace of the average value.
